# Dynamic DNA structure states interact with the RNA editing enzyme ADAR1 to modulate fear extinction memory

**DOI:** 10.1101/641209

**Authors:** Paul R. Marshall, Qiongyi Zhao, Xiang Li, Wei Wei, Abi Malathi, Esmi Zajaczkowski, Laura Leighton, Sachithrani Madugalle, Dean Basic, Ziqi Wang, Jiayu Yin, Wei-Siang Liau, Carl Walkley, Timothy W. Bredy

## Abstract

RNA modification has recently emerged as an important mechanism underlying gene diversity linked to behavioral regulation. The conversion of adenosine to inosine by the ADAR family of enzymes is a particularly important RNA modification as it impacts the physiological readout of protein-coding genes. However, not all variants of ADAR appear to act solely on RNA. ADAR1 binds directly to DNA when it is in a non-canonical, left handed, “Z” conformation, but little is known about the functional relevance of this interaction. Here we report that ADAR1 binds to Z-DNA in an activity-dependent manner and that fear extinction learning leads to increased ADAR1 occupancy at DNA repetitive elements, with targets adopting a Z-DNA structure at sites of ADAR1 recruitment. Knockdown of ADAR1 leads to an inability to modify a previously acquired memory trace and this is associated with a concomitant change in DNA structure and a decrease in RNA editing. These findings suggest a novel mechanism of learning-induced gene regulation whereby ADAR1 physically interacts with Z-DNA in order to mediate its effect on RNA, and both are required for memory flexibility following fear extinction learning.

## Introduction

RNA modification has emerged as an important aspect of RNA metabolism that has a profound influence on the function of single RNA molecules. RNA editing, the post-transcriptional conversion of single nucleotides leading to functionally distinct protein products, is one such RNA modification that has recently attracted interest for its potential role in cognition and memory^1–3^. In mammals, three different RNA editing enzymes (ADAR1-3) have been identified. ADAR1 and ADAR2 convert adenosine to inosine (A-to-I editing), where inosine is read as guanosine during transcription. A-to-I editing is known to have important downstream physiological effects. For example, *Adar1* knockout is embryonically lethal in mice, although this can be rescued by deletion of the double stranded RNA immune sensor melanoma differentiation-associated protein 5 (Mda5)^4,5^. Across the transcriptome, A-to-I editing is concentrated in short- and long-interspersed nuclear elements (SINE/LINE)^6^, which represent a population of repetitive elements that are thought to require ADAR1 mediated editing in order to alter their secondary structure and prevent the activation of Mda5^7^. In addition, ADAR1 mutations in humans is related to the development of Aicardi– Goutières syndrome, a rare neurodegenerative disease associated with cognitive dysfunction^3^. ADAR2 has both RNA binding and RNA editing domains and is best known for its role in the, editing of the ionotropic glutamate receptor 2 (GluR2) subunit^8^, which is essential for neuronal plasticity^9^. Finally, ADAR3 has an RNA binding domain and, although lacking in enzymatic activity^10^, recent work indicates that Adar3 knockout leads to impaired fear memory in mice by altering a small percentage of known RNA editing targets^11^.

Together with its RNA binding domain, ADAR1 also possesses a Z-alpha domain that enables binding to DNA when it is in a specific structural configuration, known as Z-DNA^12,13^. This DNA structure deviates from classical models of DNA, termed B-DNA^14,15^, by forming a left hand turn of the double helix and expanding the distance and electrostatic charge between the nucleotide bases^16^. In addition, early work suggested that Z-DNA maintains some rudimentary chemical memory of the previous structure occurrence^17^, which impacts the threshold for gene activation. In fact, the formation and stability of this structure depends on changes in salt^18^ and calcium^19^ concentrations, as well as DNA modification^20–24^. All processes known to occur under physiological conditions, and that are required for systems-level memory formation^25,26^. Importantly, ADAR1 directly influences active transcription, and can do so without its RNA editing domain^27^.

These observations led us to question whether the equilibrium between B-DNA and Z-DNA states may in fact influence the threshold for experience-dependent transcriptional activity^26,28^, which itself is critical for at least the first stage of memory consolidation^28^. Given that ADAR1 recognizes Z-DNA *in vitro*^29^, and potentially alters the transition from the B- to Z-DNA^30,31^, we hypothesized that ADAR1-mediated changes in DNA structure states are critical for learning-induced gene expression in the adult brain.

## Results

### Adar1 binds directly to DNA in response to neural activity, in vitro

To begin to explore the relationship between Adar1 and DNA structure states in a dynamic system, primary cortical neurons (PCNs) were stimulated and Adar1 transcript and protein levels, as well as binding to DNA were measured. *Adar1* mRNA expression increased in response to KCl-induced depolarization in PCNs (Supplementary Figure 1a). This change in gene expression was associated with a rapid, but transient, increase in Adar1 protein levels in the nucleus (Supplementary Figure 1b-c). In addition, at this time point there was an increase in Adar1 occupancy at a variety of gene targets (Supplementary Figure 1d-e and genome wide data).

### Adar1 is induced in activated neurons during fear extinction learning

*Adar1* mRNA was rapidly and transiently increased in the adult brain in response to fear extinction training (Supplementary Figure 1f). Extinction-trained mice also exhibited significantly higher levels of *Adar1* mRNA expression in activated neurons than in quiescent neurons from the same animal (Supplementary Figure 1g-h). This increased expression was selective for Adar1 in the PFC, as the expression of other known readers of Z-DNA, including Braca1 and ZBP1, were not affected (Supplementary Figure 1i-k).

### Increased Adar1 occupancy at DNA repetitive elements during fear extinction learning

To elucidate the potential relevance of experience-dependent DNA binding by Adar1, we next performed ChIP-seq on both quiescent and activated neurons following fear extinction learning. After high stringency filtering of reads and statistical correction for multiple comparisons, we found that Adar1 was significantly enriched at 122 genomic loci. When the distribution between the retention control and extinction-trained animals with high or low Arc levels was compared (Figure 1a), a significant enrichment at repetitive elements, including SINEs and LINEs proximal to peaks of Adar1 deposition in all groups was revealed (Figure 1b-c). In an independent cohort of animals that had been infused with either a scrambled control or *Adar1* shRNA into the infralimbic PFC we validated 9 targets: Neurexin 3 (Nrxn3), Carbonic anhydrase 5a, mitochondrial (Car5a), integrator complex subunit 2 (Ints2), Synaptosomal-associated protein 47 (Snap47), calcium voltage-gated channel auxiliary subunit alpha 2 delta 2 (Cacna2d2), Eukaryotic translation initiation factor 4 gamma 3 (Eig4g3), Regulator of chromosome condensation (Rcc1), sidekick cell adhesion molecule 1 (Sdk1) and glutamate ionotropic receptor kainate type subunit 2 (Grik2). All targets were selected from the genome-wide based on large statistical differences and relevance to neuronal plasticity. For example, Neurexin 3 has been shown to be critical for observational fear learning ^32^.

**Figure 1.**
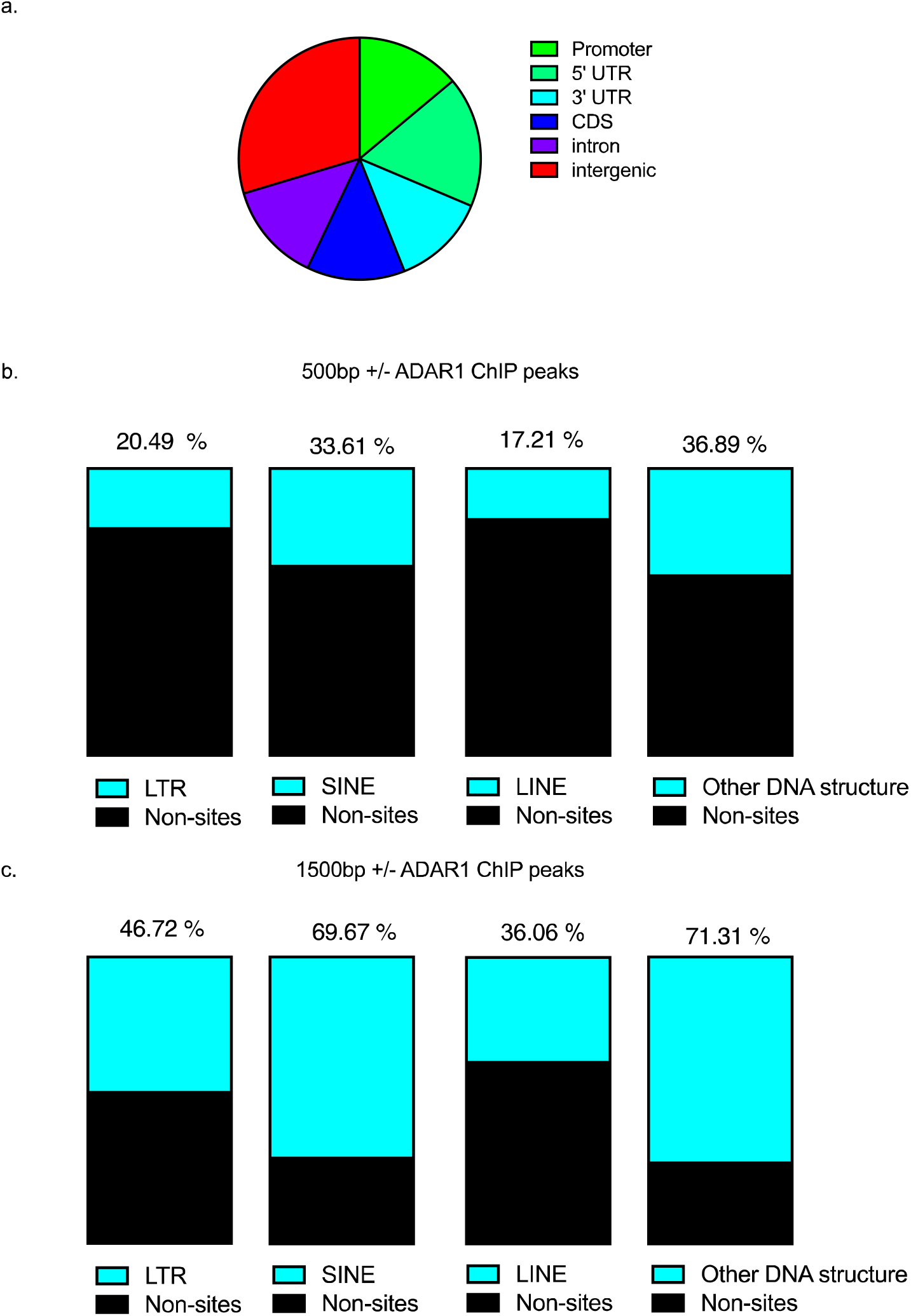
ADAR1 binds to DNA and targets DNA repetitive elements in reposnse to behavioural training. Experience-dependent distribution of Adar1 binding within ILPFC neurons of mice that were either subjected to fear conditioning training followed by exposure to a novel context (retention control: RC) or fear conditioning followed by 10 presentations of the conditioned stimuli (EXT) **a**, Mapping of Adar1 peaks from the ChIP-seq revealed that the distribution was not equally across the genome with a bias toward intergenic regions. USC genome browser annotated DNA repetitive elements **b**, 500 base pairs +/− and **c**, 1500 base pairs +/− to the identified Adar1 bioinformatic peaks.

In control animals, all targets showed a general increase in Adar1 binding following extinction training with 10 (10CS EXT) and 30 conditioned stimulus exposures (30CS EXT) (Supplementary Figure 2), but this effect was blunted following Adar1 knockdown (Supplementary Figure 3). Using structural prediction software (https://nonb-abcc.ncifcrf.gov/apps/site/default) 70% of sites that showed Adar1 enrichment had a high probability of forming alternative DNA structures, with 11% harboring known RNA editing sites (http://rnaedit.com/). These findings suggest a potential link between RNA editing and the recognition of dynamic DNA structure states at SINE/LINEs.

### Adar1 knockdown impairs memory flexibility

To assess the functional consequence of Adar1 binding to DNA, we infused an *Adar1* shRNA into the infralimbic PFC (Supplementary Figure 4a) and validated its effect on both *Adar1* mRNA and protein levels (Supplementary Figure 4b-c). In animals trained in the presence of *Adar1* shRNA (Figure 2a) there was no effect on within-trial extinction learning (Supplementary Figure 4d). Control animals that were infused with a scrambled control virus and exposed to context B (RC SC), exhibited significantly higher freezing scores at tests 1 and 2 compared to extinction trained animals treated with scrambled control (FC-EXT SC). In contrast, there was no difference in animals treated with *Adar1* shRNA and extinction trained (EXT ADAR1 shRNA) relative to RC SC mice (Figure 2b-d). Together, these data suggest three possibilities; 1) Adar1 may serve to enhance the consolidation of the original fear, 2) Adar1 may block active destabilization of the original fear memory, or 3) Adar1 may be acting more passively by stabilizing the original fear and enhancing stimulus independent memory destabilization across time.

**Figure 2.**
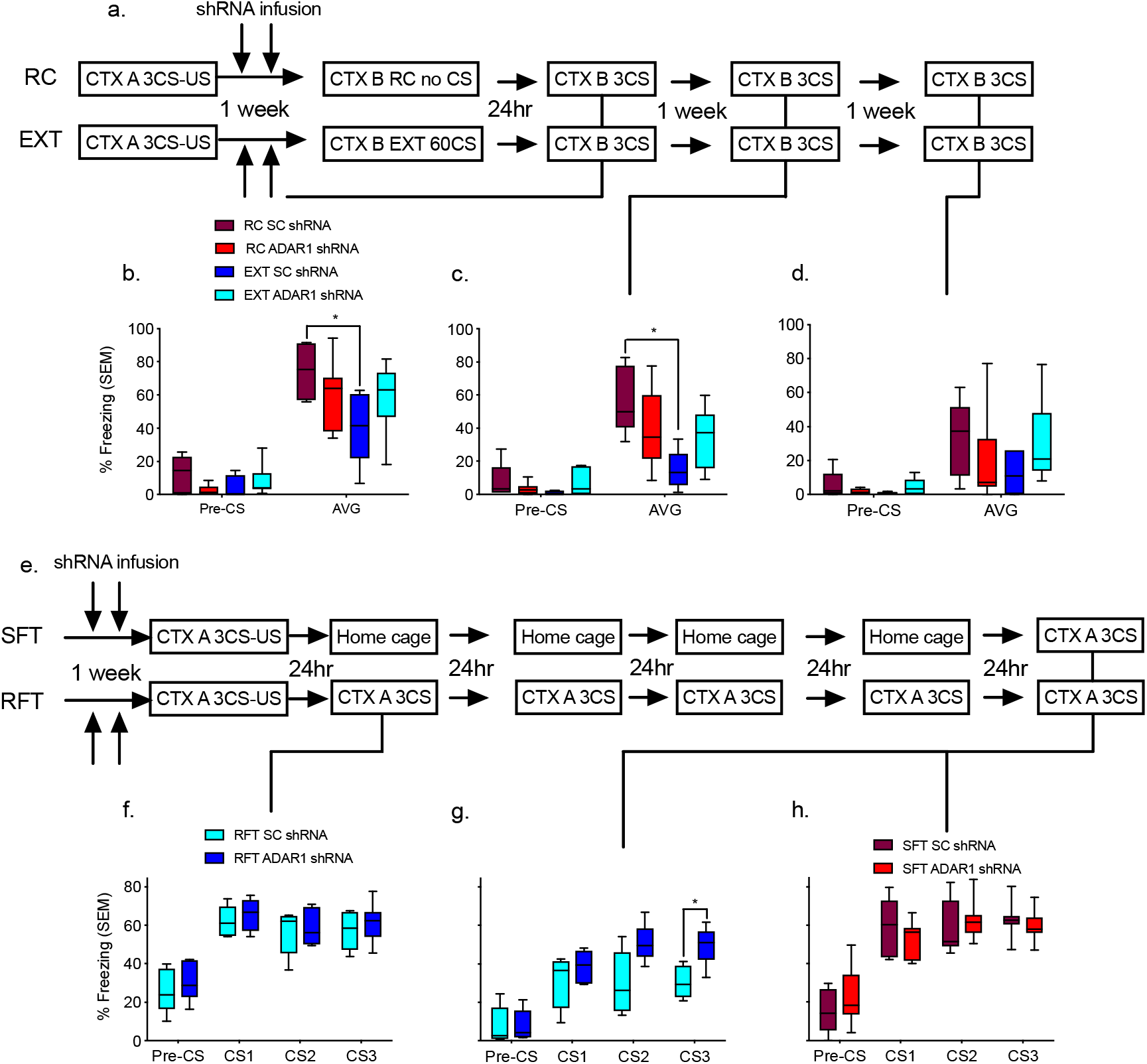
Adar1 knockdown impairs memory updating following extinction training. **a**, Schematic of the behavioural protocol used to test the effect of lentiviral-mediated knockdown of Adar1 in the ILPFC on fear extinction memory. CTX, context; CS, conditioned stimulus; US, unconditioned stimulus. **b**, Animals treated with scrambled control virus, fear conditioned and exposed to a novel context (RC SC shRNA) had significantly higher freezing scores than scrambled control animals that were fear conditioned followed by extinction (EXT SC shRNA), but this was blocked in animals that were treated with ADAR1 shRNA and extinction trained (EXT ADAR1 shRNA; n=5 biologically independent animals per group, repeated two-way ANOVA, F_1,21_ = 67.17, *p<.0001; Dunnett’s posthoc tests: RC SC vs. EXT SC, * p = 0.0462, RC SC vs EXT ADAR1 p = 0.3369, RC SC vs RC ADAR1 p = 0.4407. **c**, This effect held at the second (repeated two-way ANOVA, F_1,21_ = 85.21, *p<.0001; Dunnett’s posthoc tests: RC SC vs. EXT SC, * p = 0.0065. RC SC vs EXT ADAR1 p = 0.1568. RC SC vs RC ADAR1 p = 0.2560), **d**, but not third test (two-way ANOVA, F_1,21_ =27.56, *p<.0001; Dunnett’s posthoc tests: RC SC vs. EXT, * p = 0.0404, RC SC vs EXT ADAR1 p = 0.9994, RC SC vs RC ADAR1 p = 0.6229). **e**, Schematic of the behavioural protocol used to test the effect of lentiviral-mediated knockdown of ADAR1 on memory stability. **f**, There was no significant difference between SC and ADAR1 shRNA animals subjected to repeated fear tests (RFT) on the first day of testing **g**, however, at day 5 there was a significant difference between SC and ADAR1 shRNA (ANOVA, F_1,9_ = 5.571 *p<0.05; Bonferonni posthoc test: SC vs ADAR1 CS3 *p = 0.0326). There was also no significant difference between **h**, SC and ADAR1 shRNA animals subjected to a single fear test (SFT) on day 5.

To clarify which mechanism accounts for the effect of Adar1 knockdown, a new cohort of animals was infused with either SC or Adar1 shRNA, and then subjected to 3CS-US fear conditioning followed by either **5 days of repeated 3 CS tests in context A (RFT), or a single fear test (SFT) in context A on the 5^th^ day** (Figure 2e). SC and ADAR1 treated animals from both RFT and SFT groups did not differ during fear acquisition (Supplementary Figure 4f-h). There was also no significant difference between RFT SC and RFT ADAR1 shRNA animals in fear expression at 24hrs post-test, suggesting that ADAR1 knockdown did not alter either acquisition or consolidation of the original fear (Figure 2f). However, RFT SC showed a significant reduction in freezing on the 5^th^ test (Figure 2g) while there was also no difference between SFT SC and SFT ADAR1 shRNA at the same timepoint (Figure 2h). These data indicate that ADAR1 knockdown led to disruption in active destabilization of the original fear memory resulting in impaired extinction memory.

### ADAR1 is a critical for updating DNA structure states

To clarify the activity-dependent nature of the capacity of ADAR1 to bind to Z-DNA, two global assays were employed. One tested the effect of ADAR1 binding to a defined construct that had inserts which differed in their probability to form Z-DNA, ^33^ as this structure is known to attract ADAR1, ^12,13,34^ and our *in vitro* ChIP-seq data revealed putative targets that form this DNA structure. ADAR1 bound significantly more to GC repeats that enhanced Z-DNA (Supplementary Figure 5a-b). In the second assay, a dot blot with a Z-DNA specific antibody ^35^, using DEPC as a negative control ^35^ and spermidine treated DNA as a positive control ^36^, confirmed the presence of Z-DNA (Supplementary Figure 5c). Next, we examined the effect of neural activity on Z-DNA formation in primary cortical neurons. KCl-induced depolarization led to an increase in Z-DNA (Figure 3a). Furthermore, if an increase in Z-DNA is enhanced by spermidine or blocked by doxorubicin there would be increase or decrease, respectively, in ADAR1 binding indicating that ADAR1 negatively regulates Z-DNA (Supplementary Figure 5d-e). In line with this interpretation, we found an inverse relationship between ADAR1 and Z-DNA during extinction learning (Figure 3b). This relationship also held at genomic loci derived from the ADAR1 ChIP-seq dataset. Specifically, there were comparable levels of Z-DNA at RC and 10CS EXT, which then significantly decreased following 30CS or 60CS training (Figure 3g-j). Z-DNA levels also significantly increased immediately following fear conditioning (1hr), but not later (2-5hrs) (Figure 3c-f), and at this time there was very little ADAR1 binding and no significant groups differences (Supplementary Figure 2 h-i). Together these findings suggest that the induction of Z-DNA is required for fear learning, and its removal depends on ADAR1, which is required to modify the original trace (Figure 3g).

**Figure 3.**
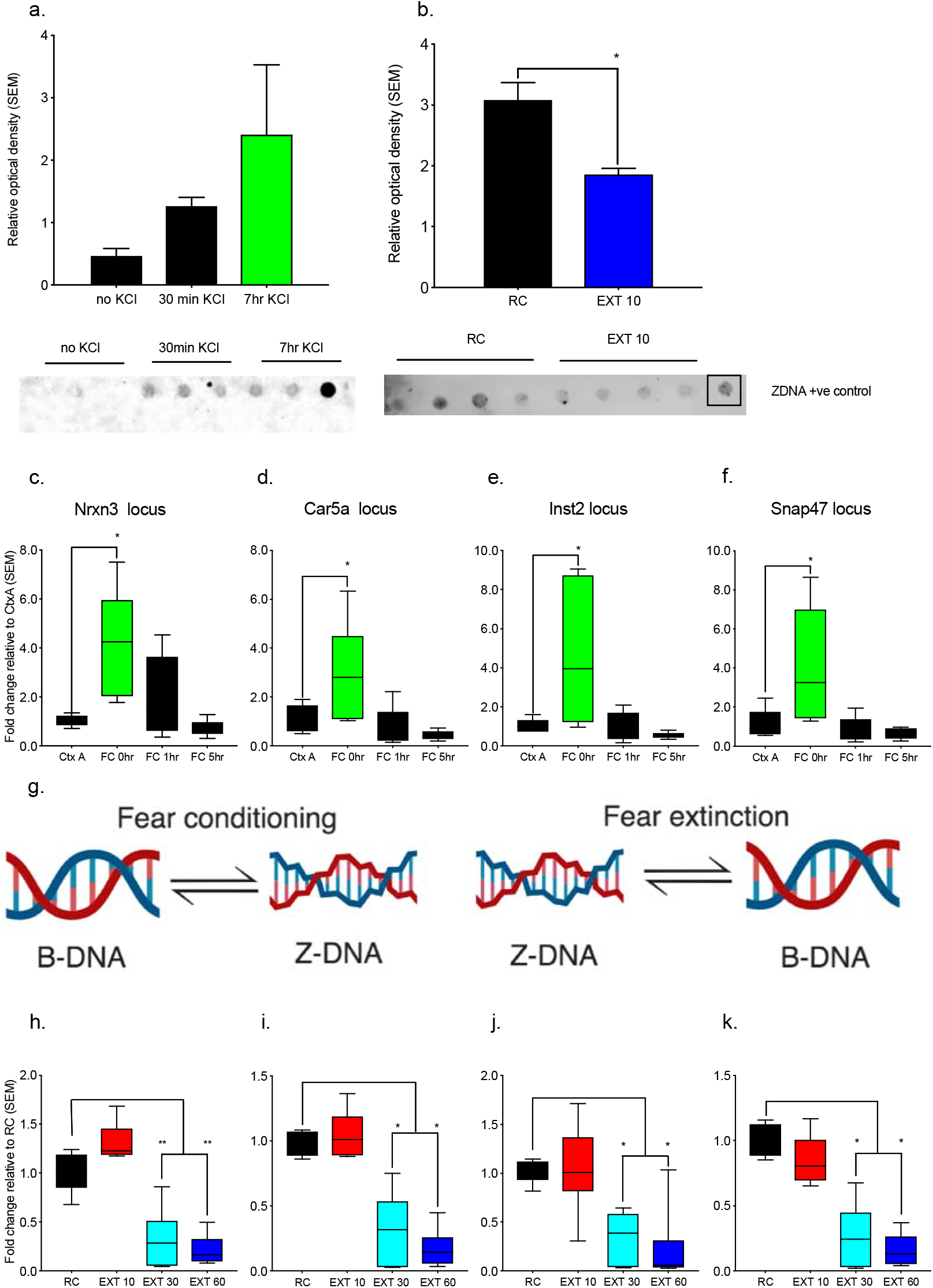
Z-DNA is a critical modulator of activity-dependent transcription. **a**, Stimulation of cortical neurons (7DIV, 20mM KCl, from 0 to 7hrs significantly altered Z-DNA structure (Kruskal-Wallis test = 5.6, *p = 0.0500). **b**, Mice exposed to fear conditioning training followed by exposer to a novel context (retention control: RC) compared to fear conditioning followed by extinction (EXT 10) had significantly lower Z-DNA t = 4.061, d.f. = 6 **p < 0.01. Animals that were exposed to the fear conditioning context but not conditioned (Ctx A) compared to those fear conditioned and euthanized immediately after training (FC O hr), 1 hr after (FC 1hr), or 5 hr after (FC 5hr) significantly increased Z-DNA at the **c**, Nrxn3 locus (ANOVA, F_3,18_ = 5.983, *p<0.01 Dunnett’s posthoc tests: Ctx A vs FC 0hr p = 0.0083, Ctx A vs FC 1hr p = 0.4994, Ctx A vs FC 5hr 0.9694), **d**, Car5a locus (ANOVA, F_3,18_ = 5.419, *p<0.01 Dunnett’s posthoc tests: Ctx A vs FC 0hr p = 0.0344, Ctx A vs FC 1hr p = 0.9087, Ctx A vs FC 5hr p = 0.6681) **e**, Inst2 (ANOVA, F_3,18_ = 5.459, *p<0.01; Dunnett’s posthoc tests: Ctx A vs FC 0hr p = 0.0142, Ctx A vs FC 1hr p = 0.999, Ctx A vs FC 5hr p = 0.9496), and **f**, Snap47 locus (ANOVA, F_3,18_ = 5.214, *p<0.01, Dunnett’s posthoc tests: Ctx A vs FC 0hr p = 0.0209, Ctx A vs FC 1hr p = 0.9807, Ctx A vs FC 5hr p = 0.9316). **g**, Model depicting the transition from B-DNA to Z-DNA during fear conditioning and from Z-DNA to B-DNA during fear extinction. There was also a significant alteration in Z-DNA when comparing fear trained animals exposed to a novel context following fear conditioning (retention control: RC) to fear conditioned followed by 10CS (EXT 10), 30 CS (EXT 30), or 60CS extinction (EXT 60). Specifically, Z-DNA significantly decreased at the **h**, Nrxn3 locus (ANOVA F_3,23_ = 39.48 ****p<0.0001, Dunnett’s posthoc tests: RC vs EXT 10 p = 0.0879 RC vs EXT 30 ****p <0.0001, RC vs EXT 60 <0.0001), **i**, Car5a locus (ANOVA F(_3,22_) = 32.94 ****p<0.0001, Dunnett’s posthoc tests: RC vs EXT 10 p = 0.9031 RC vs EXT 30 ****p <0.0001, RC vs EXT 60 ****p<0.0001), **j**, Inst2 locus (ANOVA, F_3,25_ = 13.88, ****p<0.0001, Dunnett’s posthoc tests: RC vs EXT 10 p = 0.9920, RC vs EXT 30 p = 0.0009, RC vs EXT 60 p = 0.0002), and **k**, Snap47 locus (ANOVA, F_3,23_ = 34.88, ****p<0.0001, Dunnett’s posthoc tests: RC vs EXT 10 p = 0.3363, RC vs EXT 30 ****p < 0.0001, RC vs EXT 60 ****p<0.0001).

### ADAR1 disinhibits Z-DNA thereby promoting flexible transcription

To provide further support for the hypothesis that DNA structure states are important for learning, we next explored how ADAR1 binding and Z-DNA modulation may be functionally linked through transcription. The expression of mRNA proximal to intergenic DNA targets known to be involved in extinction learning, such as Nrxn3, mirrored the profile of Z-DNA. For both fear conditioning and extinction, Z-DNA formation outside of protein coding regions was significantly correlated with the expression of the protein coding genes (see Table 1). In addition, there was an increase in RNA polymerase II (RNA polII) occupancy during extinction at all candidates, and a significant reduction during fear training, which was mirrored by a significant increase in Z-DNA (Supplementary Figure 6a-h). With respect to Nrxn3, Z-DNA initially increased, which was followed by a significant increase in ADAR1 binding, leading to a reduction in Z-DNA and increased expression of the upstream transcript (Figure 4 a-d).

**Figure 4.**
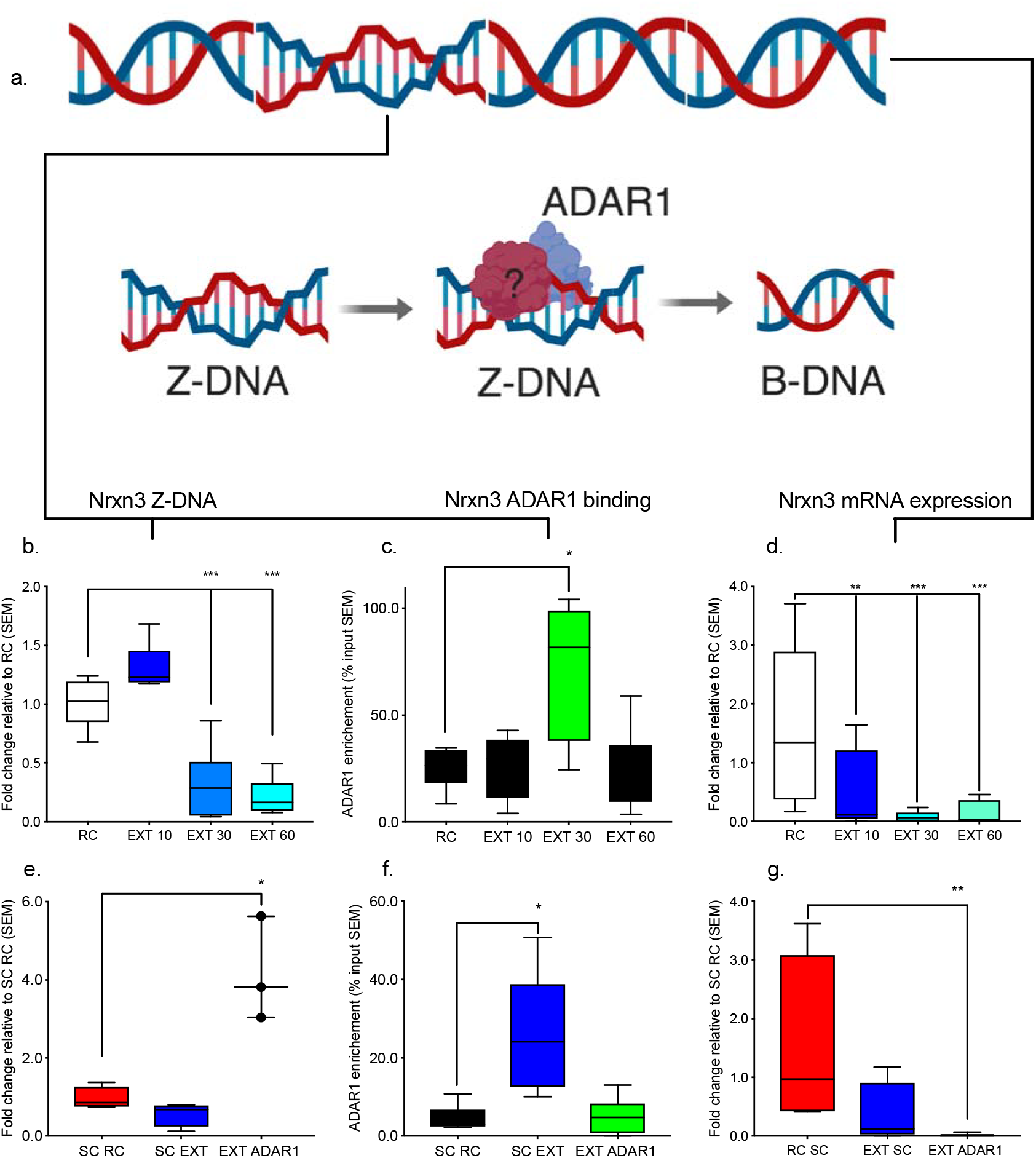
Adar1-Z-DNA interaction at Nrxn3 locus modulates the expression of transcripts derived from a non-coding region enriched with SINEs. **a**, Model depicting the recognition and resolution of Z-DNA by Adar1 binding **b**, There was a significant reduction in Z-DNA when comparing fear trained animals exposed to a novel context following fear conditioning (retention control: RC) to fear conditioned followed by 10CS (EXT 10), 30 CS (EXT 30), or 60CS extinction (EXT 60) for the Nrxn3 locus (ANOVA F_3,23_ = 39.48 ****p<0.0001, Dunnett’s posthoc tests: RC vs EXT 10 p = 0.0879 RC vs EXT 30 ****p <0.0001, RC vs EXT 60 <0.0001). Further there was a significant increase in **c**, Adar1 at the 30CS EXT time point (ANOVA F_3,23_ = 8.130 ***p<0.001, Dunnett’s posthoc tests: RC vs EXT 10 p = 0.9993 RC vs EXT 30 ****p <0.0001, RC vs EXT 60 p > 0.9999). **d**, In response to extinction training, Nrxn3 transcript was significantly decreased (ANOVA, F_3,18_ = 4.636 *p<0.01, Dunnett’s posthoc tests: RC vs EXT 10 p = 0.0938 RC vs EXT 30 **p = 0.0066, RC vs EXT 60 p 0.0251), **e**, When these tests were conducted with ADAR1 shRNA treated animals Z-DNA was now significantly increased (ANOVA, F_2,8_ = 24.96 ***p<0.001, Dunnett’s posthoc tests: SC RC vs SC EXT 10 p = 0.6690 SC RC vs ADAR1 EXT ***p = 0.0007), **f**, ADAR1 shRNA also blocked the significant increase in ADAR1 binding (ANOVA, F_2,15_ = 9.798 **p<0.01, Dunnett’s posthoc tests: SC RC vs SC EXT 10 **p = 0.0027 SC RC vs ADAR1 EXT p = 0.9897), **g**, Further ADAR1 shRNA treated animals had significantly lower Nrxn3 transcript, (ANOVA, F_2,10_ = 3.345 p = 0.0772, Fishers LSD posthoc tests: SC RC vs SC EXT 10 **p = 0.0956 SC RC vs ADAR1 EXT *p = 0.0307).

**Table 1.** Relationship between Z-DNA and transcription. Correlations are between the amount of Z-DNA enrichment at Adar1 binding sites and transcript levels of exon expression from targets proximal to the sites.

| Gene target | Group |  |  |  |
| --- | --- | --- | --- | --- |
|  | Fear conditioned |  | Extinction trained |  |
|  | Correlation | P-value | Correlation | P-value |
| Car5a | 0.5671 | 0.0176 | -0.3522 | 0.1391 |
| Snap47 | 0.2885 | 0.2614 | -0.4733 | 0.0261 |
| Inst2 | 0.2368 | 0.0559 | 0.4632 | 0.0820 |
| Nrxn3 | -0.1343 | 0.6073 | 0.3582 | 0.1444 |

ADAR1 knockdown had the opposite effect and led to a significant increase in Z-DNA, reduced ADAR1 binding, and a further reduction in Nrxn3 RNA expression (Figure 4 e-g). Z-DNA may therefore serve as a negative regulator of transcription at these sites, which is normally disinhibited by ADAR1. Genomic updating by dynamic changes in Z-DNA though ADAR1 binding may therefore be a critical mechanism underlying gene expression associated with memory flexibility.

### RNA editing at SINEs and LINEs require ADAR1 binding to DNA

Because ADAR1 is also critically involved in RNA editing we assessed whether ADAR1 binding at non-coding sites also impacted RNA editing. Since these sites were proximal to annotated SINE/LINEs (https://genome.ucsc.edu/) we first established whether there was RNA expression directly by exploring raw reads from a previously published RNA sequencing experiment (Li et al, 2019; Data not shown). We then designed primers around these regions, which overlapped with ADAR1 targets, and performed PCR. Gel electrophoresis followed by sequencing confirmed RN expression at overlapping ADAR1 binding sites. Using bioinformatic prediction software (http://sines.eimb.ru/), several putative SINE/LINEs were detected within the ADAR1 genome-wide data set and confirmed. Furthermore, the expression of these elements was significantly altered by both fear conditioning and fear extinction learning (Figure 5a-e). We next performed polymerase III (PolIII) ChIP at the same sites and found that there was significant increase in PolIII binding at the Nrxn3 locus following extinction. Although ADAR1 knockdown impaired this effect, a reintroduction of full length ADAR1 led to a complete rescue of the effect of fear extinction learning on Nrxn3 mRNA expression (Supplementary Figure 6i-l).

**Figure 5.**
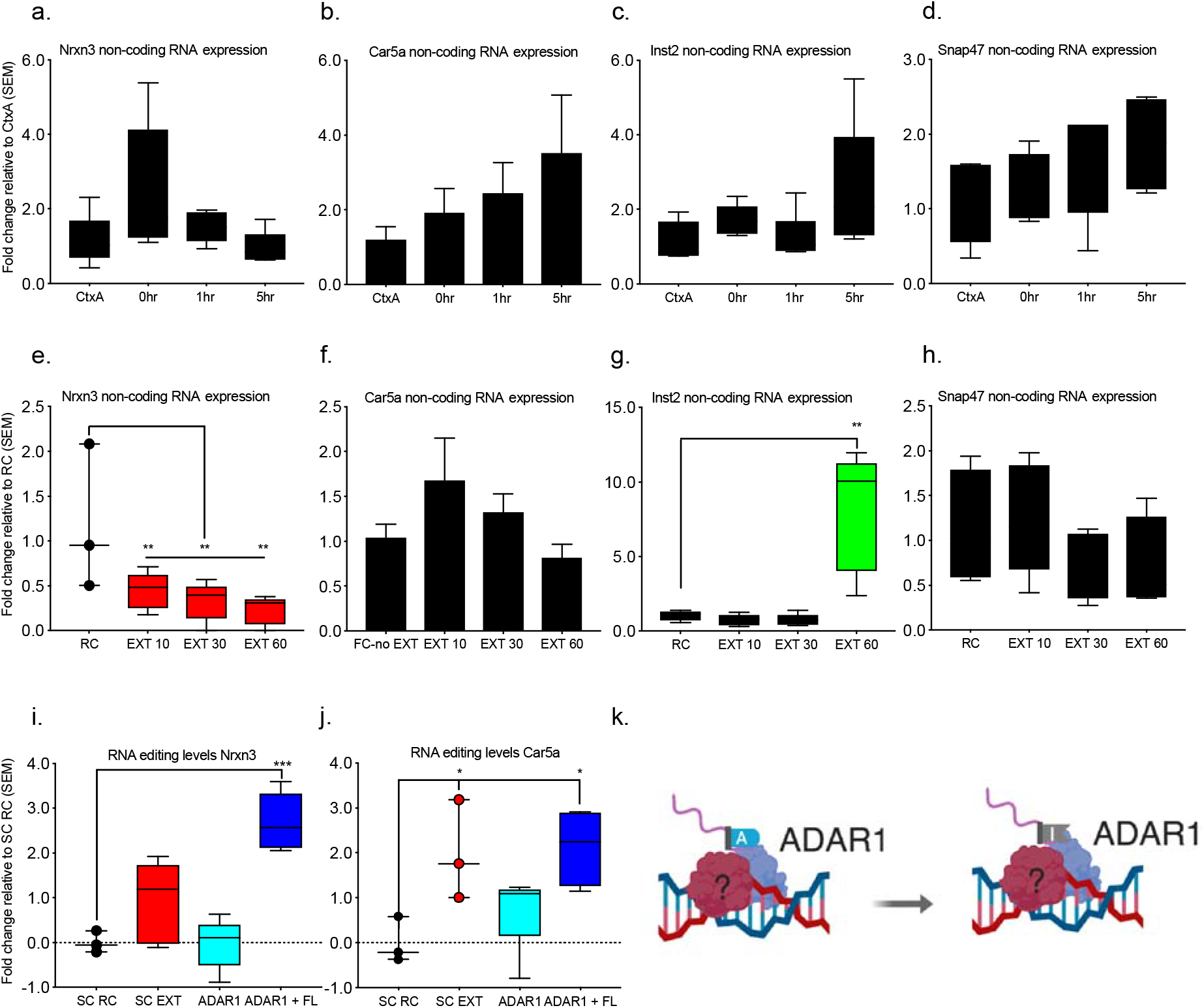
Editing of RNA derived from SINEs and LINEs requires Adar1 binding to DNA. RNA expression at the sites of Adar1 binding outside of the exons did not reach statistical significance for animals that were exposed to the fear conditioning context but not conditioned (Ctx A) compared to those fear conditioned and euthanized immediately after training (FC O hr), 1 hr after (FC 1hr), or 5 hr after (FC 5hr) in the **a**, Nrxn locus (ANOVA, F_3,17_ = 0.0946), **b**, Car5a locus (ANOVA, F_3,14_ = 0.3320), **c**, Inst2 locus (ANOVA, F_3,16_ = 0.1590), or **d**, Snap47 locus (ANOVA, F_3,13_ = 0.3756). Fear trained animals exposed to a novel context following fear conditioning (retention control: RC) compared to fear conditioned followed by 10CS (EXT 10), 30 CS (EXT 30), or 60 CS extinction training (EXT 60) instead showed a significant reduction of expression at the **e**, Nrxn3 locus (ANOVA, F_3,14_ = 5.016, *p < 0.05. Dunnett’s posthoc tests: RC vs EXT 10 p = 0.0342, RC vs EXT 30 p = 0.0145, RC vs EXT 60 p = 0.0069). As well as a significant increase in RNA expression at the **g**, Inst2 locus (ANOVA, F_3,15_ = 15.15, ****p < 0.0001. Dunnett’s posthoc tests: RC vs EXT 10 p = 0.9925, RC vs EXT 30 p = 0.9924, RC vs EXT 60 p = 0.0004). However, the **f**, Car5a (ANOVA, F_3,13_ = 1.812, p = 0.1946) and **h**, Snap47 locus (ANOVA, F(_3,15_) = 1.233, p = 0.3326) did not reach statistical significance. EXT trained animals injected with scrambled control shRNA (EXT SC), ADAR1 shRNA (EXT ADAR1), or ADAR1 shRNA and full length ADAR1 (EXT ADAR1 + FL) were compared to animals subject to fear conditioning and injected with scrambled control shRNA (RC SC) and assessed for RNA editing with Endo V treatment. Significant increases in RNA editing levels were observed at the Nrxn3 locus **i**, (ANOVA, F_3,14_ = 16.72, ****p < 0.0001. Dunnett’s posthoc tests: SC RC vs SC EXT p = 0.1913, SC RC vs ADAR1 EXT, p = 0.9999, SC RC vs ADAR1 + FL EXT, ***p = 0.0002), as well as the **j**, Car5a locus (ANOVA, F_3,11_ = 4.687, *p<0.05 Dunnett’s posthoc tests: SC RC vs SC EXT p = 0.0424, SC RC vs ADAR1 EXT, p = 0.5063, SC RC vs ADAR1 + FL EXT, *p = 0.0206). **k**, Pictorial representation of ADAR1 enzymatically converting adenosine to inosine.

Next, in order to assess whether changes in ADAR1 binding to DNA were associated with RNA editing, we used Endo V, an enzyme that recognizes and cuts inosine in RNA followed by qPCR. In support of the hypothesis that an interaction between Z-DNA and ADAR1 may be associated with co-transcriptional RNA editing, there was significant increase in RNA editing in response to fear extinction learning, and this effect was blocked by ADAR1 knockdown. Importantly, simultaneous introduction of full length ADAR1 rescued the observed effect of ADAR1 knockdown on Z-DNA and RNA editing (Figure 5 g-k). Together these findings strongly suggest a dual function of ADAR1.

### Memory flexibility requires both ADAR1 Z-DNA binding and RNA editing domains

To determine whether the observed impairment in fear extincion is dependent on both ADAR1 DNA binding and RNA editing, three variants of ADAR1 were introduced into the infralimbic PFC post fear training (Figure 6 c). Either full length ADAR1, ADAR1 with a mutated z-alpha domain (N175A/Y179A) which blocks its ability to bind DNA, or ADAR1 with a mutated RNA editing domain (E861A) that blocks its ability to RNA edit, was introduced against an ADAR1 knockdown background and compared to SC animals subject to either RC or EXT conditions (Figure 6 a-b). At the first test, all groups that been exposed to extinction training showed lower freezing scores than control animals except those treated with ADAR1 shRNA (Figure 6 c-f). On the second and third tests, only the SC EXT and ADAR1 + Full length EXT group had significantly lower freezing scores. These findings. indicate that the z-alpha and RNA editing domain are both required to rescue memory flexibility.

**Figure 6.**
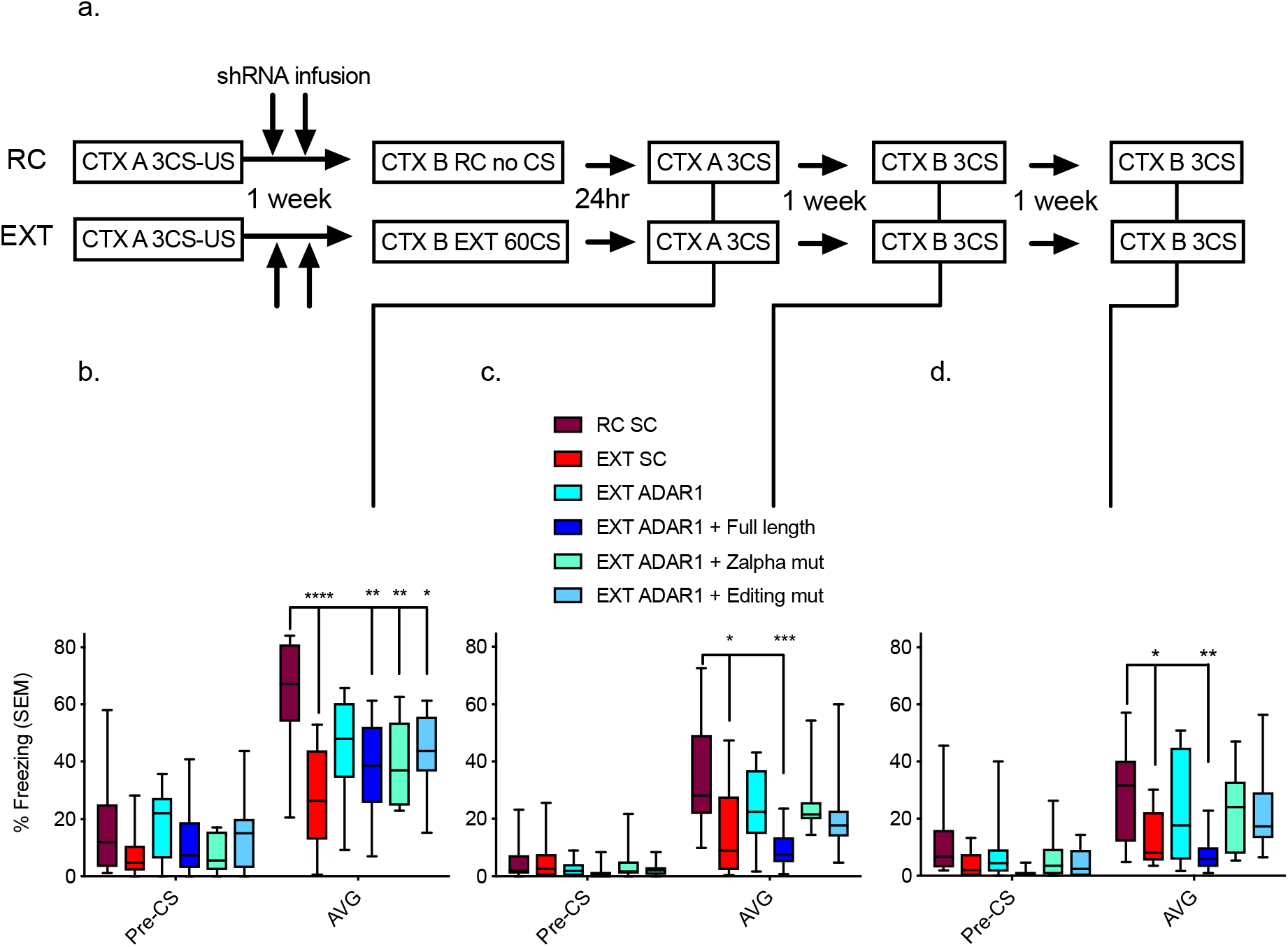
Fear extinction requires both Adar1 Z-DNA binding and RNA editing domains. **a**, Schematic of the behavioural protocol used to test the effect of lentiviral-mediated knockdown of ADAR1 and restoration using either ADAR1 + FL, ADAR1 + Zalpha mutant, or ADAR1 + Editing mutant, on extinction learning. CTX, context; CS, conditioned stimulus; US, unconditioned stimulus. **b**, Animals treated with scrambled control virus, fear conditioned and exposed to a novel context (SC RC) had significantly higher freezing scores than scrambled control animals that were fear conditioned followed by extinction (SC EXT), but this was blocked in animals that were treated with any of the other three viruses (n=11 biologically independent animals per group, repeated two-way ANOVA, F_5,60_ = 5.353, ***p<0.001; Dunnet post-hoc; SC RC vs. SC EXT, **** p < 0.0001, SC RC vs ADAR1 EXT p = 0.0697, SC RC vs ADAR1 + FL EXT **p = 0.0022, ADAR1 + Z-alpha mut **p = 0.0044, ADAR1 + Edit mut *p = 0.0334). **c**, During the second test only the SC EXT and ADAR1 + FL EXT were significantly different from SC RC (n=11 biologically independent animals per group, repeated two-way ANOVA, F_5,60_ = 3.861 **p<0.01; Dunnet post-hoc; SC RC vs. SC EXT, * p = 0.0192, SC RC vs ADAR1 EXT p = 0.3487, SC RC vs ADAR1 + FL EXT ***p = 0.0009, ADAR1 + Z-alpha mut p = 0.4766, ADAR1 + Edit mut p = 0.2542). **d**, Similarly during the third test only the SC EXT and ADAR1 + FL EXT groups were significantly different from SC RC (n=11 biologically independent animals per group, repeated two-way ANOVA, F_5,60_ = 2.736, *p<0.05; Dunnet post-hoc; SC RC vs. SC EXT, *p = 0.0130, SC RC vs ADAR1 EXT p = 0.4897, SC RC vs ADAR1 + FL EXT **p = 0.0019, ADAR1 + Z-alpha mut p = 0.3247, ADAR1 + Edit mut p = 0.3153).

## Discussion

We have discovered that the formation of Z-DNA occurs during fear conditioning and, subsequently, this mark maintains an inverse relationship with gene expression, and coordinates editing of transcripts derived from putative SINE/LINE elements during fear extinction learning. We have found that ADAR1 interacts with over 100 genomic regions in activated neurons in response to fear extinction learning, and that this triggers a rapid change in Z-DNA structure, as well as an increase in PolII occupancy and gene expression, which is followed by increased RNA editing. Importantly, these effects are abolished following ADAR1 knockdown. Finally, we demonstrate that that the ability of ADAR1 to modulate flexibility of fear memory is dependent on both its DNA binding and RNA editing domains.

The idea that ADAR1 could interact with DNA structure dates back to original observations made by Rich and colleagues who suggested that the Z-alpha domain of ADAR1 recognizes an alternative left-handed DNA structure ^12,13,34^. Over the years, a very strong case has been made for the existence of Z-DNA and its role in many biological processes, particularly transcriptional regulation ^37–40^. Specifically, it has been reported that Z-DNA is related to both transcriptional activation ^41,42^ and inhibition ^43,44^. Mechanistically, this change in structure is thought to be coupled to transcription by the recruitment of transcription factors, such as PolII, for enhancement, or the direct blockade of the transcriptional machinery leading to PollI stalling during inhibition ^37,39,45–47^. Similarly, we observed that Z-DNA was both positively and negatively correlated to gene expression in response to learning, with positive correlations tending to occur during fear conditioning, and negative ones occurring following extinction training in the same locus, suggesting a reversible mechanism which also dictates memory flexibility ^25^.

Our findings confirm that ADAR1 binds to Z-DNA in an inducible manner in the adult brain, and suggest that ADAR1 actively regulates this structure to modify learning-related transcription. These observations align with the role of other proteins containing Z-DNA binding domains, such as protein kinase R (PKR) and Neurofibromin 1 (NF1), in transcriptional regulation^42,48–50^. Like Z-DNA readers, it has also been shown that the binding of ADAR1 to DNA can drive it towards a Z-DNA state ^51^. However, ADAR1 binding to DNA was minimal during the formation of Z-DNA at these sites during fear suggesting ADAR1 wasn’t driving Z-DNA formation. More surprisingly, extinction-specific reductions in Z-DNA mediated by ADAR1 suggest an unknown mechanism whereby ADAR1 recruits a helicase or other DNA repair enzyme to cut and release the torsion from Z-DNA ^52,53^, as well as releasing the poised PolII, as has been observed with topoisomerase II beta (Top2b) during fear learning ^54,55^. This would also suggest that damage at regions of DNA assuming a Z-conformation ^56^ and the role of ADAR1 as a transcriptional activator ^27,49^ are, in fact, functionally linked.

In addition, we found that a reduction in ADAR1-mediated editing of previously uncharacterized RNAs derived from SINE/LINE elements impairs fear extinction, as both the editing domain and z-alpha domain were required. Restoring the Z-alpha domain without the editing domain was not as effective as restoring both suggesting that the changes in DNA structure and RNA editing are interdependent and temporally linked such that DNA structure changes occur first followed by binding and RNA editing by ADAR1 and other attracted enzymes. This also begs the question of how non-coding RNA editing might affect memory flexibility. The edited transcripts can either: facilitate further RNA editing^57^, limit activation of the MDA5 pathway^7^, or act as functional gene enhancers by coupling PolIII- to PolII-mediated transcription^58^. A limitation of the current model is that not all ADAR1 sites identified by sequencing appear to be related to Z-DNA. There may be additional DNA structures or motifs that also recruit ADAR1, such as the G-quadruplex structure, which has been shown to be related to ADAR1 in HeLa cells ^59^. Furthermore, not all genomic loci bound by ADAR1 align with annotated genes known to be subject to RNA editing.

In summary, ADAR1 appears to show both sequence and structural specificity. We have discovered that the recognition and modification of Z-DNA structure by ADAR1 is required for memory flexibility such that the removal of this domain impairs the ability of fear extinction to overcome previously established fear memories. Together, these data support prior observations that RNA modification is required for learning, and include dynamic changes in DNA structure as an important factor in the regulation of gene expression underlying flexible memory states.

## Acknowledgements

The authors gratefully acknowledge grant support from the NIH (R01MH105398-TWB) and the NHMRC (GNT1145172 and GNT1160823-TWB), the ARC (GNT190101078-XL), the Westpac Future Scholars program (ELZ, LJL and SM) and postgraduate scholarships from NSERC (PRM) and the University of Queensland (PRM, ELZ, LJL, DB, JY, and SM). We would also like to thank Ms. Rowan Tweedale for helpful editing of the manuscript.

## Methods

### Materials and Methods

#### Animals

9-11 week-old C57BL/6 male mice were housed four per cage, maintained on a 12hr light/dark schedule, and allowed free access to food and water. To allow for identification of behavioural outliers, animals were transferred to pair housed conditions and split with a plexiglass divider at least 24hrs prior to training. All testing was conducted during the light phase in red-light-illuminated testing rooms. All animal use and training, including use of embryos, followed protocols approved by the Animal Ethics Committee of the University of Queensland and in accordance with the Australian Code for the Care and Use of Animals for Scientific Purposes (8th edition, revised 2013).

#### Primary cortical neuron

Cortical tissue was isolated from embryonic day 16 embryos. Primary cortical neurons were isolated by removing the skull and meninges with fine tipped tweezers after extracting the embryos’. Cells were then mixed into a media solution containing Neurobasal medium (GIBCO 21103) containing 5% serum, B27 supplement (GIBCO 17504-044) and 0.5-1% Penicillin-Streptomycin (GIBCO 15140) and made homogenous with gentle pipetting. The cells were then passed through a 40 micrometre cell strainer (BD Falcon 352340) and plated onto 6 well cell culture dishes coated with Poly-L-Ornithine (Sigma P2533) at a density of 1 million cells per well.

#### RNA and DNA Extraction

Both cultured cells and tissue from mice was extracted and then placed in PBS. Gentle pipetting was used for *in vitro* preparations to generate a homogenous solution. Tissue was prepared by dounce homogenization in 500 μl of PBS and RNA/DNA extracted using Trizol. For RNA, the Trizol reagent was used according to the manufacturer’s instructions (Invitrogen). DNA extraction was carried out using DNeasy Blood & Tissue Kit (Qiagen) with RNAse A (5 prime), RNAse H and RNAse T1 treatment (Invitrogen). Both extraction protocols were conducted according to the manufacturer’s instructions. The concentration of DNA and RNA was measured by Qubit assay (Invitrogen).

#### qRT-PCR

1μg of RNA was used for cDNA synthesis prepared from the Quantitect Reverse Transcription Kit according to the manufactures’ protocol (Qiagen). Quantitative PCR was then performed on a RotorGeneQ (Qiagen) real-time PCR cycler with SYBRGreen Master mix (Qiagen), using primers for target genes and beta actin or phosphoglycerate kinase as an internal control. The threshold cycle for each sample was chosen from the linear range and converted to a starting quantity by interpolation from a standard curve run on the same plate for each set of primers. All mRNA levels were normalized for each well relative to internal control using the ΔΔCT method, and each PCR reaction was run in duplicate for each sample and repeated at least twice.

#### Western blot

Individual wells from cultured cells and tissue from mice was collected and then the homogenate was fractionated utilizing a NXTRACT CelLytic™ NuCLEAR™ Extraction Kit (Sigma) according to the manufacturer’s instructions. Briefly, samples were prepared on ice (to a final volume of 40 microliters) in a mixture of Laemmli Sample Buffer and 2-Mercaptoethanol (BioRad) and then vortexed and denatured for 10 min at 70°C. Gels were run and proteins transferred onto nitrocellulose membrane (Hybond-ECL, Amersham). The membrane was blocked with Odyssey Blocking Buffer (LiCore P/N 927-40100) for 1 hour at room temperature, washed with Trizma buffered saline with 1% triton (TBST) for 5 min (3x) and incubated with 5ml of mouse monoclonal antibody (ADAR1 1:1000; (Santa Cruz sc-73408 B) in blocking buffer for 24 hours at 4°C. The membrane was washed with TBS-T (3x), incubated for 1 hour with IRDye 680 rabbit anti-mouse secondary antibody (1:10000; LI-COR) in TBS-T, and washed in TBST for 10 min (3x). Optical density readings of the blots were made using the LI-COR analysis system.

#### Dot blot

The protocol was modified from Li *et al* 2019. DNA negative controls were prepared by combing DNA with 40mM DEPC, positive controls were treated with 1M spermidine after which DNA purification was performed. Following this, DNA was spotted onto nitrocellulose membranes followed by 10 minutes of incubation at room temperature. Membranes were then hybridized with 260nm UV light, and incubated with 1:1,000 dilution of Z-DNA antibody (abcam with biotin conjugated to it with the Lightning-Link (R) Rapid Type B Biotin Antibody Labelling Kit) at 4 °C overnight. After three rounds of washes with 1X TBST, the membrane was incubated with 1:20,000 goat anti-rabbit secondary antibody conjugated with 680RD Streptavidin (LiCOR). The intensity score of each dot was then analysed by the Odyssey Fc system and normalized to background. Equivalent amount of DNA was aliquoted at the beginning of the blot and subject to RT-qPCR for PGK. This value was used to normalize the optical density score to input.

#### Lentiviral delivery and behavioural analysis

Double cannula (PlasticsOne) were implanted in the anterior posterior plane, along the midline, into the infralimbic prefrontal cortex (ILPFC). The injection locations were centred at +1.8 mm in the anterior-posterior plane (AP), and −2.8 mm in the dorsal-ventral plane (DV). Animals were then separated into single housing and given at least one week to recover before being behaviourally trained. Following the first day of training a total volume of two microliters of lentivirus per hemisphere was infused via two one microliter injections over a 48hr period. Mice were first fear conditioned, followed by 2 lentiviral infusions 24 hours post-fear conditioning, and, after a one-week of incubation, the mice were either extinction trained or just exposed to context B for an equivalent period of time (see Figure 2A). In brief, this training consisted of two contexts (A and B). Both conditioning chambers (Coulbourn Instruments) had two transparent walls and two stainless steel walls with a steel grid floors (3.2 mm in diameter, 8 mm centers); however, the grid floors in context B were covered by flat white plastic transparent surface. Additionally, context A was sprayed with a dilute lemon smell, and context B was sprayed with a dilute vinegar smell to minimize context generalization. Cameras within the boxes captured movement and were processed automatically with a freezing measurement program (FreezeFrame). The training protocol consisted of 120 sec pre-fear conditioning incubation; followed by three pairing of a 120 sec, 80dB, 16,000 Hz tone (CS) co-terminating with a 1 sec (2 min intervals), 0.7 mA foot shock (US). Mice were matched into equivalent treatment groups based on freezing during the third training CS. For extinction, mice were exposed in context B in which they habituated to chamber for 2 min, and then, extinction training (EXT) comprised of 10, 30 or 60 non-reinforced 120 sec CS presentations (5-sec intervals). For the retention control (RC) animals, context exposure was performed following fear conditioning, but without presentation of the tones. For the retention tests, all mice were returned to either context A or B and following a 2 min acclimation (used to minimize context generalization), freezing was assessed during three 120sec CS presentations (120 sec intertribal interval). Memory was inferred by the percentage of time spent freezing during the tests.

#### ADAR1 and ADAR1 scrambled control (SC) knockdown lentiviral constructs

Lentiviral plasmids were generated by inserting either ADAR1 or ADAR1 SC shRNA using the following sequences or ADAR1: GATCCCCGCCGAGTCAGTGTTTATGATTTTCAAGA GAAATCATAA ACACTGACTCGGCTTTTTTC ADAR1 SC: GATCCCCAGTTCATTAG GCTAACGTATTTCAAG AGAATACGTTAGCCTAATGAACTTTTTTTC immediately downstream of the human H1 promoter in a modified FG12 vector (FG12H1, derived from the FG12 vector originally provided by David Baltimore, CalTech). Lentivirus was prepared and maintained according to previously published protocols ^61^.

#### Adar1, Adar Za mutant and catalytically inactive Adar1

murine Adar1, Adar1 Za-mutant (N175A/Y179A) and editing dead mutant (E861A) were generated by gene synthesis and mutants introduced by gBlocks (IDTDna). All plasmids were sequence verified. The cDNA was cloned into pLVX vector (Clontech).

#### Immunohistochemistry

Following the end of testing mice were euthanized with 100mg/Kg ketamine, after which 20ml of saline was pumped through the circulatory system followed by 4% paraformaldehyde to fix the tissue. The brains were then placed in 30% sucrose for a minimum 24hr prior to cryostat slicing. Sectioning at 40um was performed using a cryostat and sections were mounted on Menzel-Glaser Superfrost Plus microscope slides. Briefly, the sections were incubated 1-2hr in blocking buffer, after which primary antibodies (ADAR1 and GFP ab6556) were added and the slides incubated at 4°C overnight. The slides were then washed 3 times with PBS containing 0.02% Tween 20 (PBS-T), after which secondary antibodies were added. The slides were then incubated at room temperature for 2hr, washed 3 times with PBST, and cover-slipped with ProLong Gold Antifade Mountant with DAPI (P36931). The sections were then imaged on a slide scanning confocal microscope.

#### FACS

The procedure for sorting activated neurons for cDNA preparation and ChIP-seq was modified from published protocols (Li *et al* 2019). Briefly, following sample preparation for FACS the identified population of neurons, as indicated by their high intensity in the 488nm channel, was further split into two populations which had intensity in the 647nm Arc channel above the upper part of the non-NeuN population (High Arc) or below it (Low Arc; see Supplementary Figure 1D). 250,000 cells from each sample was then taken and four biological replicates were pooled to make one for further processing to reach at least 1 million cells required for reliable chromatin immunoprecipitation.

#### Chromatin and carrier ChIP immunoprecipitation (ChIP)

Standard ChIP was performed following modification of the Invitrogen ChIP kit protocol. Lysate from cells or tissue was fixed in 1% formaldehyde and cross-linked cell lysates were sheared by Covaris in 1% SDS lysis buffer to generate chromatin fragments with an average length of 300bp. For samples not being subjected to sequencing 1 million yeast (Kindly provided by Micheal Kobor) per sample were spiked in prior to fixation to enhance antibody-target interactions when cell count is low ^62^. Following shearing, the chromatin was then immunoprecipitated using the Adar1 Antibody preconjugated to biotin (sc-73408 B), Anti-Z DNA antibody (ab2079), Anti-RNA polymerase II CTD repeat YSPTSPS (phospho S5) antibody (ab5131), Anti-POLR3A antibody (ab96328), or normal rabbit IgG (Santa Cruz), overnight at 4°C. Protein-DNA-antibody complexes were then precipitated with Dynabeads Protein G (Thermofisher 10003D) or Dynabeads MyOne Streptavidin C1 (Thermofisher 65001) for 1hr at 4°C, followed by three washes in low salt buffer, and three washes in high salt buffer for protein G beads, and 3 washes in biotin wash buffer, followed by 2 washes in PBS. The precipitated protein-DNA complexes were eluted from the antibody with 1% SDS and 0.1M NaHCO3 and incubated for four hours at 65C in Proteinase K. Following proteinase K digestion, phenol-chloroform extraction, and ethanol precipitation, samples were subjected to qPCR using primers specific for 200bp segments corresponding to the target regions.

#### ChIP-Seq data analysis

Bioinformatic analysis performed as reported in Li *et al* 2019. After removing the duplicate reads, low mapping quality reads and paired-end reads that were not properly aligned, MACS2 (v.2.1.1.20160309) was used to call peaks for each sample using the parameter setting ‘callpeak -t SAMPLE -c INPUT -f BAMPE–keep-dup□=□all -g mm -p 0.05 -B’. Peak summits identified by MACS2 from all samples were collected to generate a list of potential binding sites. Custom PERL script was then applied to parse the number of fragments (hereafter referred to as counts) that covered the peak summit in each sample. Each pair of properly paired-end aligned reads covering the peak summit represented one count. The total counts in each sample were normalized to 20 million before comparison among samples. The potential binding sites were kept if they met all of the following conditions: (1) the sites were not located in the mm10 empirical blacklists and (2) the normalized counts in all three biological replicates in one group were larger than in its input sample, and the normalized counts in at least two replicates were more than twofold larger than in the normalized input count.

#### RNA editing assay

Following extraction of RNA, 500ng per sample was either combined with 10 units of Endo V (NEB M0305S) in 1X NEBbuffer 4 (NEB #B7004), or an equivalent volume of ultra-pure water and 1X NEBbuffer 4. According to manufacture protocol, both reactions were incubated at 37C for 1hr, followed by 65C for 5 minutes to inactivate. RNA was further purified by Zymo RNA clean concentrator kit and then amplified with the RT enzyme, buffer, and random primer mix from Qiagen (Cat No. 205313) at 42C for 30 minutes, and 95C for 3 minutes. Following this cDNA samples were subjected to standard qPCR comparing Endo V treated to untreated for each target.

