## Supplementary Figures 1-7 for "Dynamic DNA structure states interact with the RNA editing enzyme ADAR1 to modulate fear extinction memory"

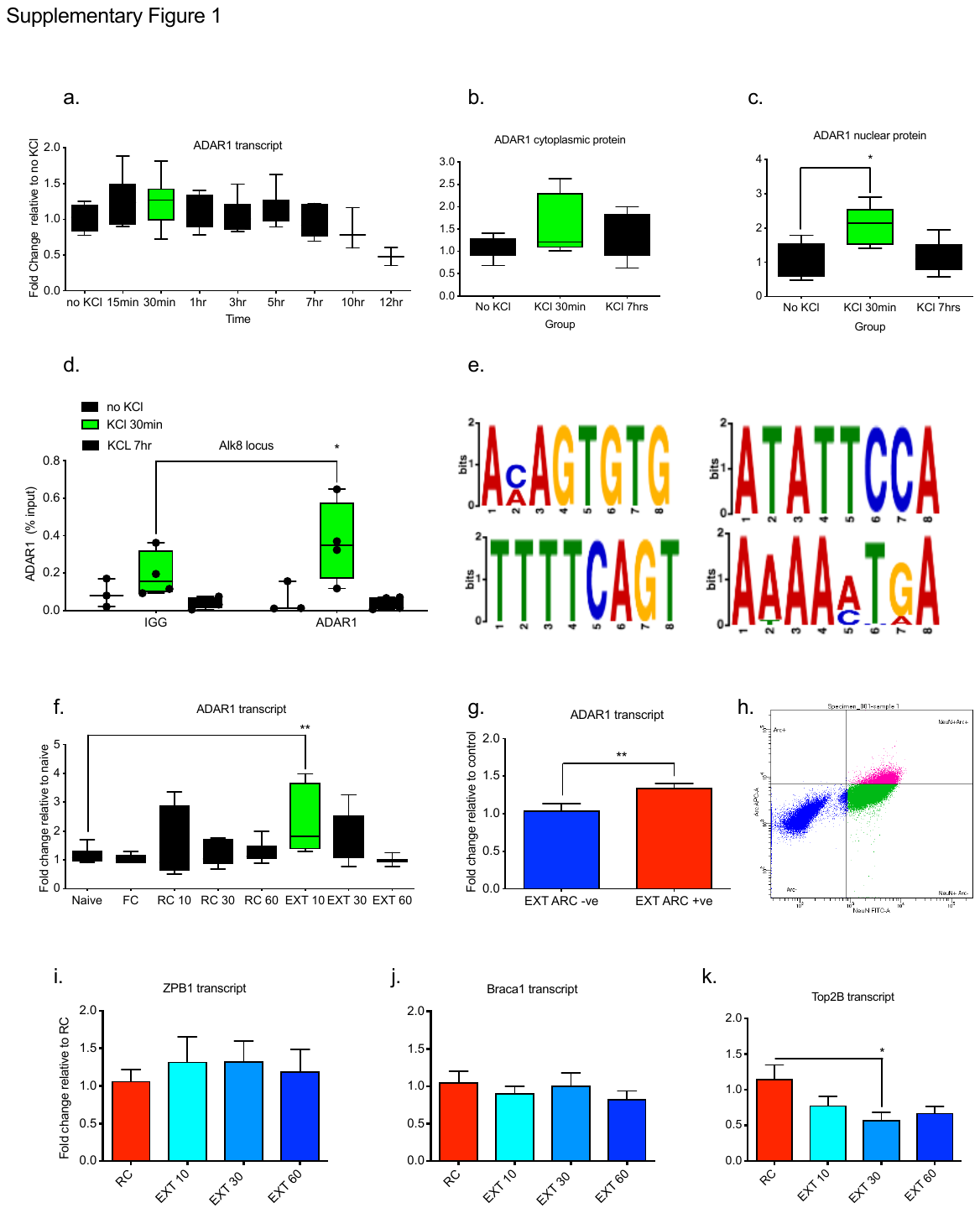

**Supplementary Figure 1. ADAR1 binds directly to DNA in response to neural activity, in vitro. a**, Stimulation of cortical neurons (7DIV, 20mM KCl, from 0 to 12 hrs) significantly altered ADAR1 transcript levels (ANOVA F_8,48_ = 2.78, p<0.05 **c,** and significantly enhanced nuclear ADAR1 protein levels (ANOVA F_2,12_ = 5.63, *p =.019. Dunnet post-hoc at 30 minutes post-stimulation *p=0.017) **b,** but did not significantly alter cytoplasmic ADAR1 protein levels, (ANOVA F_2,12_= 1.1, p = 0.36). **d,** Stimulation also significantly increased ADAR1 at the Alk8 locus Two-way ANOVA, Main effect of KCl (0, 30min or 7hr F_2,8_ = 5.14, *p=0.0367), Main effect of antibody (IGG vs ADAR1 F_1,8_ =4.95, p=0.056 and interaction F_2,8_ = 9.52, **p=0.007 (2,16) = 3.93; Post hoc comparison between IGG and ADAR1 at 30 minutes t = 3.18 d.f. = 3 *p=0.0251), **e,** Using the MEME-ChIP program against all bioinformatically significant targets ADAR1 DNA binding motifs were identified. **f,** Naïve home-cage control mice compared to those fear trained (FC), exposed to a novel context following fear conditioning (retention control: RC) or fear conditioned followed by 10CS (EXT 10) 30 CS (EXT 30) or 60CS extinction (EXT 60) led to a significant enhancement of ADAR1 transcript (ANOVA, F_5,39_ = 3.49, p = 0.005, Dunnett post-hoc relative to Naïve, 30CS was significantly increased *p=0.011). **g,** RNA transcript levels for EXT 10 animals which are either: high in the neuronal marker NeuN, and activity-regulated cytoskeleton-associated protein (EXT ARC +ve) or high in the neuronal marker NeuN and low in ARC (EXT ARC -ve) are significantly different t = 2.706, d.f. = 6, *p<0.05 **h,** Representative fluorescence-activated cell sorting plot showing the separation between neurons and non-neurons based on NeuN, and the gating of Arc levels into high and low. RNA transcript levels from mice fear trained by 3CS-US pairing followed by exposure to context B with no sounds (RC), compared to animals fear trained and exposed to context B with 10 (EXT 10) 30 (EXT 30) or 60 (EXT 60) presentations of the tone that was previously paired with the shock, there was no significant alteration in **i,** Z-DNA-binding protein 1 (ZBP1) and **j,** Breast cancer type 1 susceptibility protein (Braca1), **k,** although there was a significant decrease for DNA Topoisomerase II Beta (Top2B) (ANOVA, F_3,26_ = 3.157, *p<0.05 Dunnet post hoc; RC vs EXT 30 *p = 0.0229).

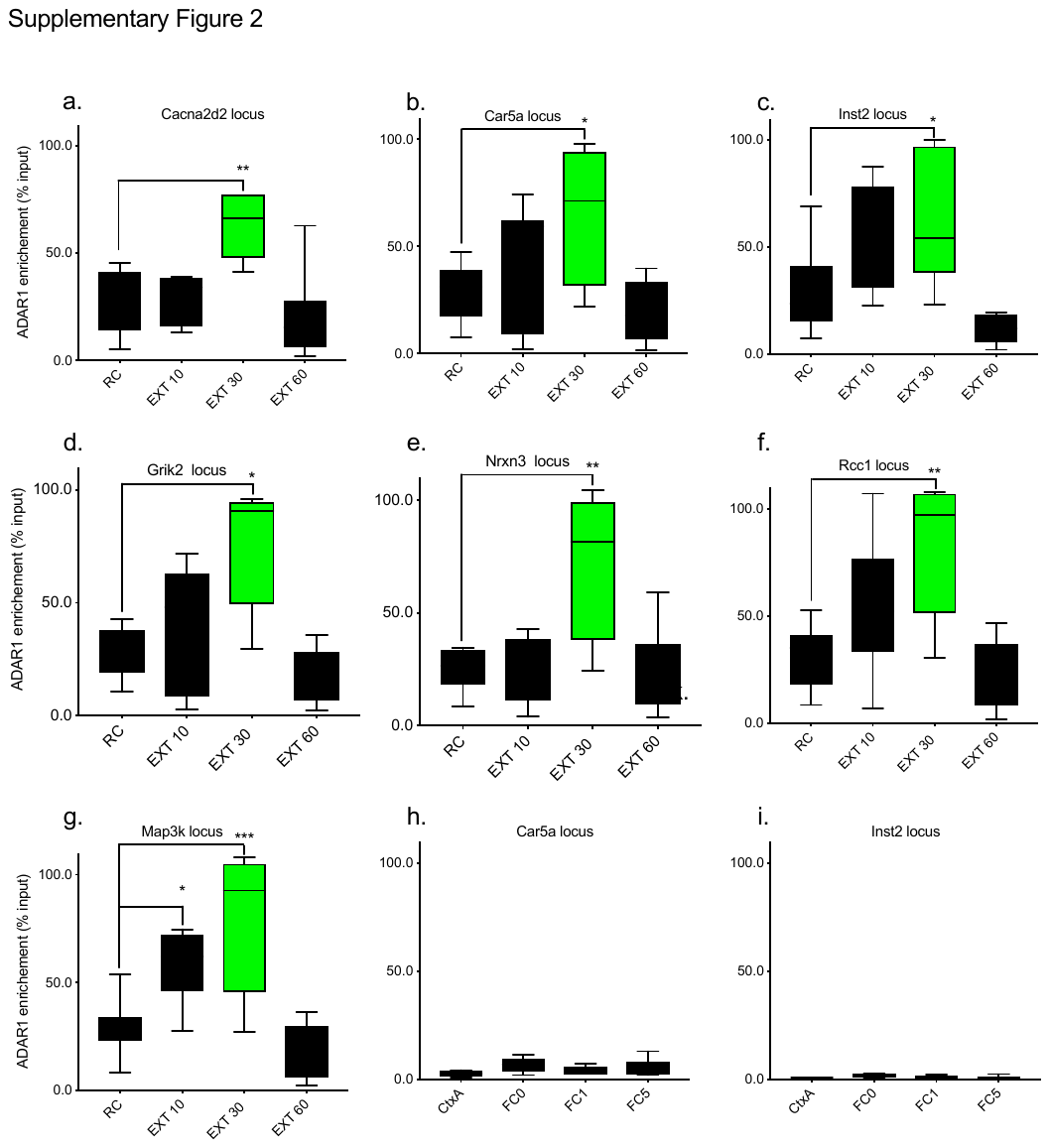

**Supplementary Figure 2. ADAR1 occupancy across the genome.** Animals exposed to a novel context following 3CS-US fear conditioning (retention control: RC) were compared to those exposed to 10CS (EXT 10) 30 CS (EXT 30) or 60CS extinction (EXT 60) all targets showed a significant increase in ADAR1 enrichment at the 30CS time point this includes **a,** Cacna2d2 locus (ANOVA, F_3,22_ = 8.107 ***p<0.001, Dunnett’s posthoc tests: RC vs EXT 10 p = 0.9733 RC vs EXT 30 **p = 0.0021, RC vs EXT 60 p = 0.7828 **b,** Car5a locus (ANOVA, F_3,21_ = 4.517 *p<0.05, Dunnett’s posthoc tests: RC vs EXT 10 p = 0.4417 RC vs EXT 30 *p = 0.0475, RC vs EXT 60 p = 0.7429), **c,** Inst2 locus (ANOVA, F_3,23_ = 8.599 ***p<0.001, Dunnett’s posthoc tests: RC vs EXT 10 p = 0.0736 RC vs EXT 30 *p = 0.0169, RC vs EXT 60 p = 0.2972), **d,** Grik2 locus (ANOVA, F_3,22_ = 9.050 ***p<0.001, Dunnett’s posthoc tests: RC vs EXT 10 p = 0.5679 RC vs EXT 30 *p = 0.0023, RC vs EXT 60 p = 0.6247), **e,**  Nrxn3 locus (ANOVA, F_3,23_ = 8.130 ***p<0.001, Dunnett’s posthoc tests: RC vs EXT 10 p = 0.9993 RC vs EXT 30 **p = 0.0016, RC vs EXT 60 p = 0.999), **f,** Rcc1 locus (ANOVA, F_3,23_ = 7.677 ***p<0.001, Dunnett’s posthoc tests: RC vs EXT 10 p = 0.2177 RC vs EXT 30 **p = 0.0037, RC vs EXT 60 p = 0.7799), **g,** Map3k (ANOVA, F_3,22_ = 13.08 ****p<0.001, Dunnett’s posthoc tests: RC vs EXT 10 *p = 0.0252 RC vs EXT 30 ***p = 0.0005, RC vs EXT 60 p = 0.6002). However targets **h,** Car5a and **i,** Inst2 both did not show any significant changes follow fear conditioning at any of the time points.

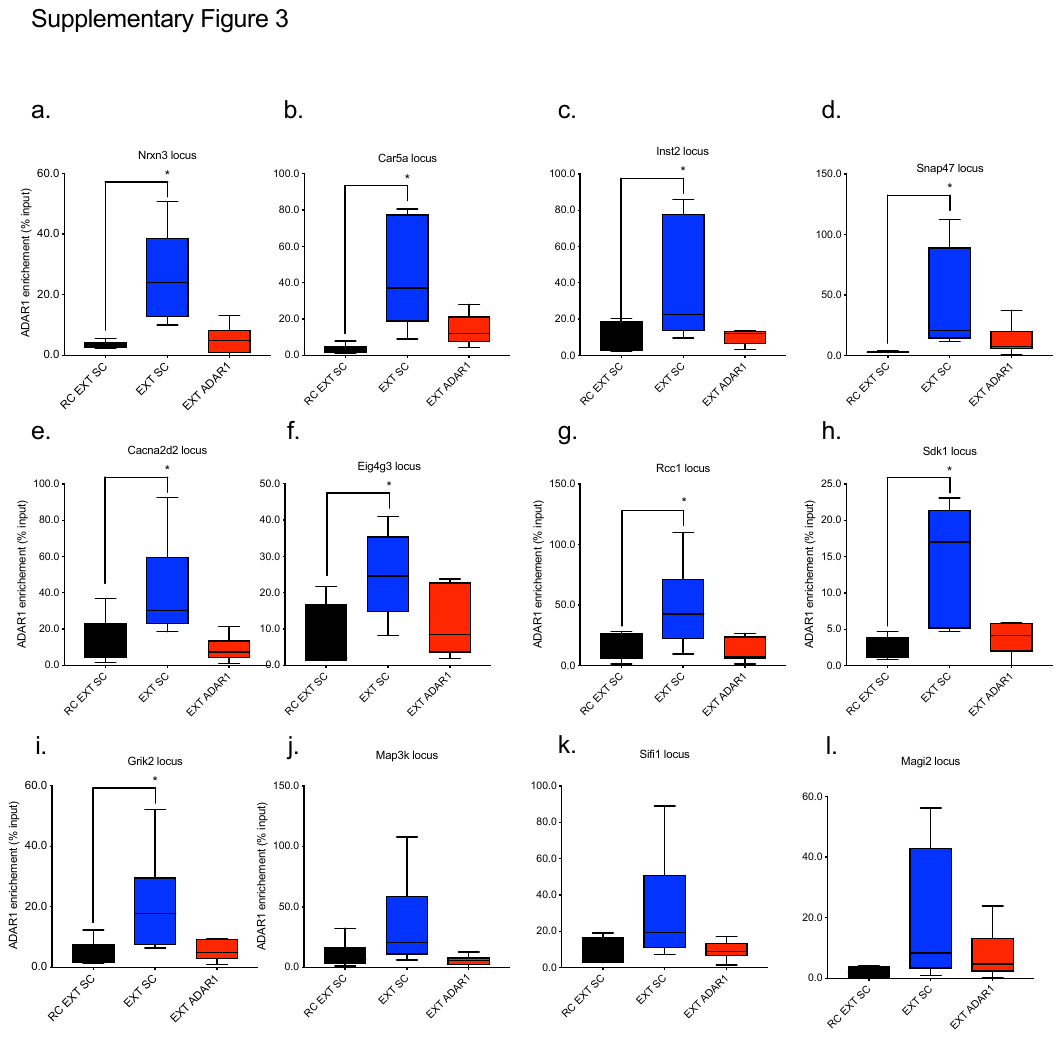

**Supplementary Figure 3. ADAR1 shRNA validation.** Animals exposed to a novel context following 3CS-US fear conditioning (retention control: RC) were compared to those exposed to 10CS EXT and infused with either scramble control shRNA (EXT SC) or 10CS EXT and ADAR1 shRNA (EXT ADAR1). For all 12 targets tested there was an increase for extinction which was blocked by ADAR1 shRNA significantly at the following loci: **a,** Nrxn3 (ANOVA, F_2,14_ = 9.831 **p<0.01, Dunnett’s posthoc tests: SC RC vs SC EXT 10 **p = 0.0029 SC RC vs ADAR1 EXT p = 0.9121), (b) Car5a (ANOVA, F_2,14_ = 6.356 *p<0.01, Dunnett’s posthoc tests: SC RC vs SC EXT 10 **p = 0.0083 SC RC vs ADAR1 EXT p = 0.5904), **c,** Inst2 (ANOVA, F_2,16_ = 4.494 *p<0.05, Dunnett’s posthoc tests: SC RC vs SC EXT 10 *p = 0.0359 SC RC vs ADAR1 EXT p = 0.9979), **d,** Snap47 (ANOVA, F_2,16_ = 4.760 *p<0.05, Dunnett’s posthoc tests: SC RC vs SC EXT 10 *p = 0.0277 SC RC vs ADAR1 EXT p = 0.7363), **e,** Cacna2d2 (ANOVA, F_2,14_ = 9.831 **p<0.01, Dunnett’s posthoc tests: SC RC vs SC EXT 10 **p = 0.0029 SC RC vs ADAR1 EXT p = 0.9121), **f,** Eig4g3 (ANOVA, F_2,14_ = 4.378 *p<0.05, Dunnett’s posthoc tests: SC RC vs SC EXT 10 *p = 0.0284 SC RC vs ADAR1 EXT p = 0.7952), **g,** Rcc1 (ANOVA, F_2,26_ = 9.577 ***p<0.001, Dunnett’s posthoc tests: SC RC vs SC EXT 10 **p = 0.0021 SC RC vs ADAR1 EXT p = 0.9353), **h,** Sdk1 (ANOVA, F_2,11_ = 6.637 *p<0.05, Dunnett’s posthoc tests: SC RC vs SC EXT 10 *p = 0.0138 SC RC vs ADAR1 EXT p = 0.8611), **i,** Grik2 (ANOVA, F_2,14_ = 4.415 *p<0.05, Dunnett’s posthoc tests: SC RC vs SC EXT 10 *p = 0.0380 SC RC vs ADAR1 EXT p = 0.9696). While **j,** Map3k **k,** Sifi1 and **l,** Magi2 loci did show trends in a similar direction they did not reach significance.

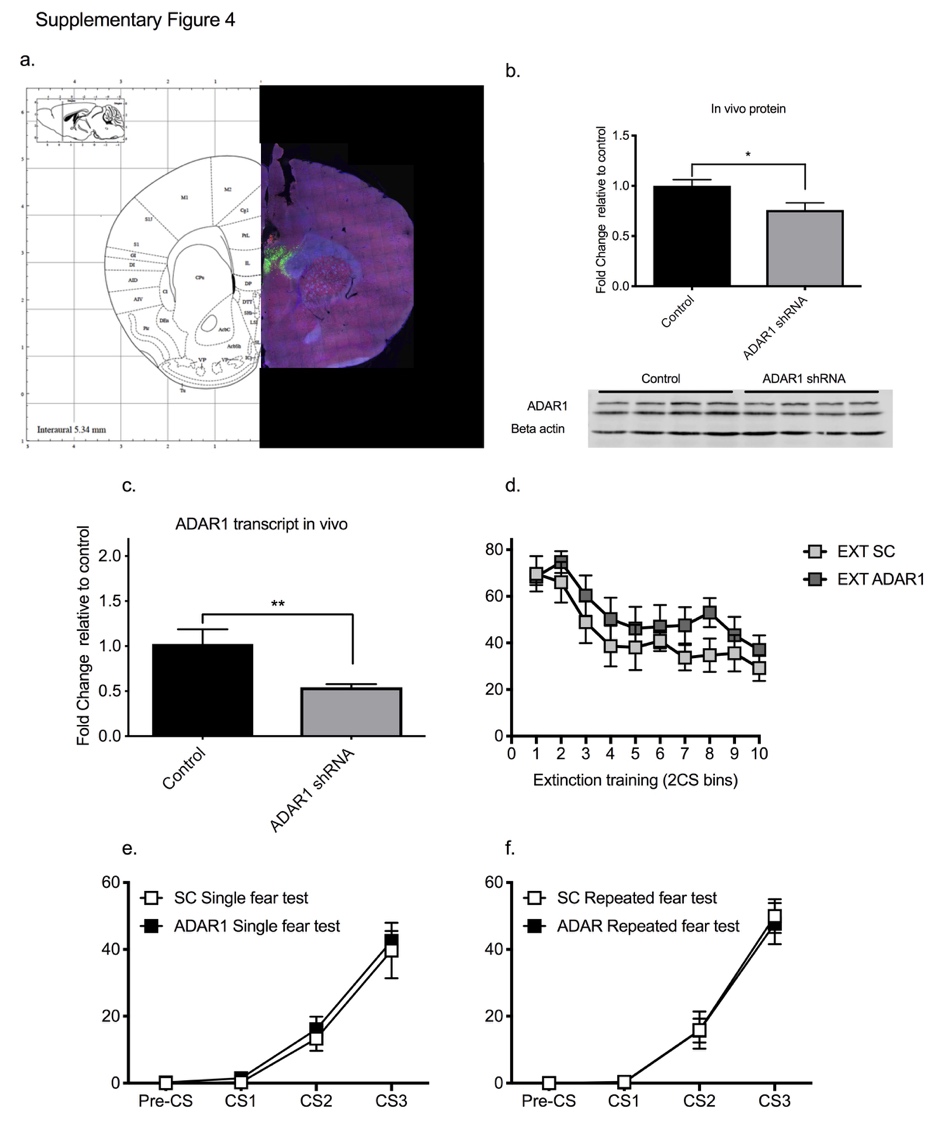

**Supplementary Figure 4. Behavioral validation for interpretation a,** Representative image of cannula placement in the ILPFC, DAPI in blue, ADAR1 staining in red, GFP from ADAR1 shRNA transfection in green. **b,** ADAR1 shRNA significantly reduced ADAR1 protein levels t = 2.54, d.f. = 6 *p<0.05. **c,** ADAR1 shRNA also significantly reduced ADAR1 transcript relative to animals treated with a scrambled control virus t = 2.891, d.f. = 8 *p<0.05. **d,** There was no significant difference in freezing levels during extinction training for animals treated with SC shRNA and ADAR1 shRNA. Also no significant difference in freezing scores during fear acquisition for animals treated with SC or ADAR1 shRNA and subject to either **e,** repeated (RFT) or **f,** a single fear test (SFT).

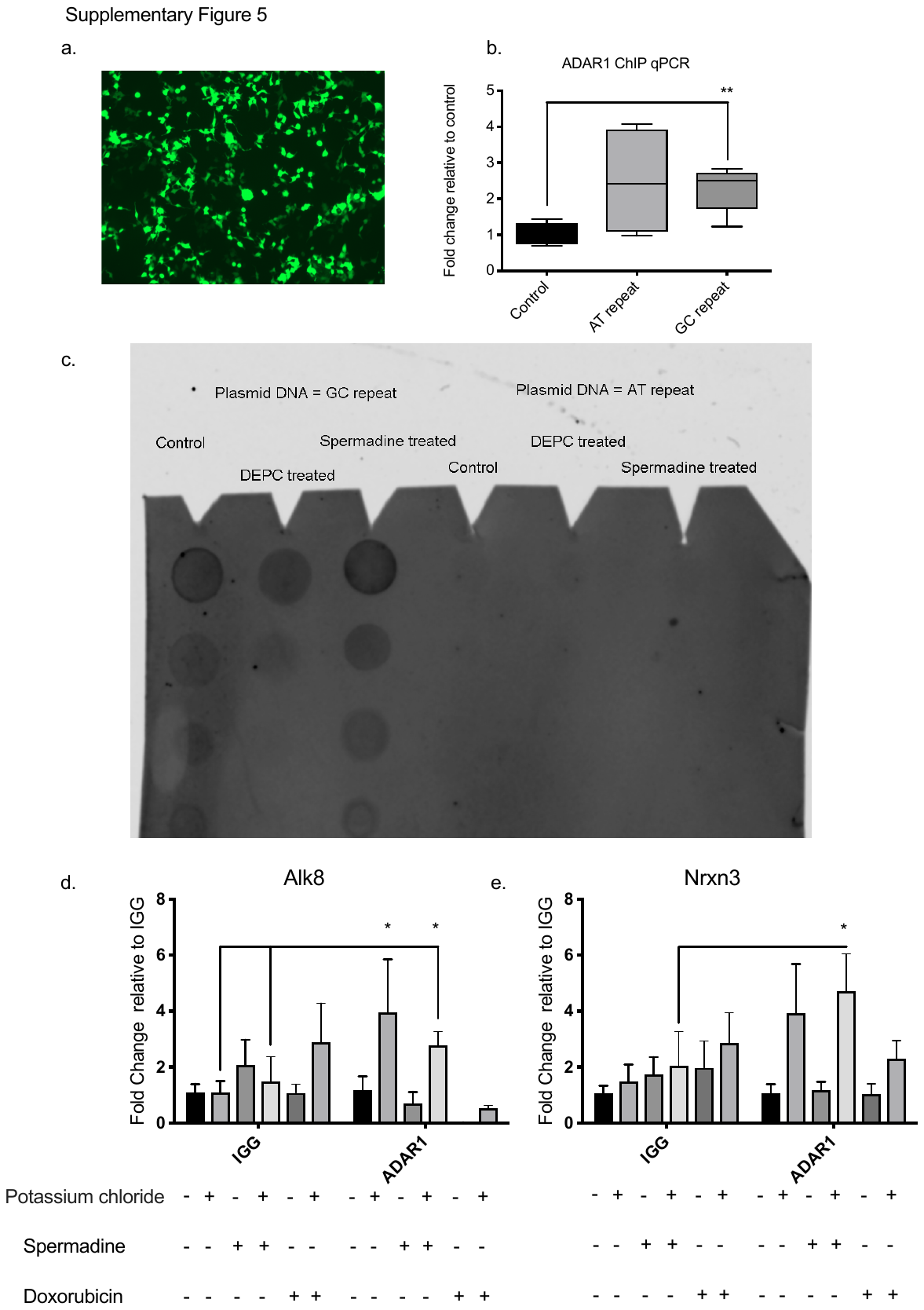

**Supplementary** **Figure 5**. **Validation of Z-DNA measurements**

**a**, Representative image of transfection of FG12 construct containing GFP reporter and either a control, a AT rich, or GC rich sequence. **b**, There is a significant enrichment of ADAR1 binding to constructs containing GC repeats. **c,** Blots demonstrating that Z-DNA antibody can detect positive controls containing GC repeats and this signal is increased by 1M spermidine and blocked by 40mM DEPC. **d**, Stimulation of primary cortical neurons with 10mM KCl led to significant enrichment of ADAR1 at the Alk8 locus for those treated with control and spermidine. For Nrxn3 only cells treated with spermidine led to a significant enrichment of ADAR1.

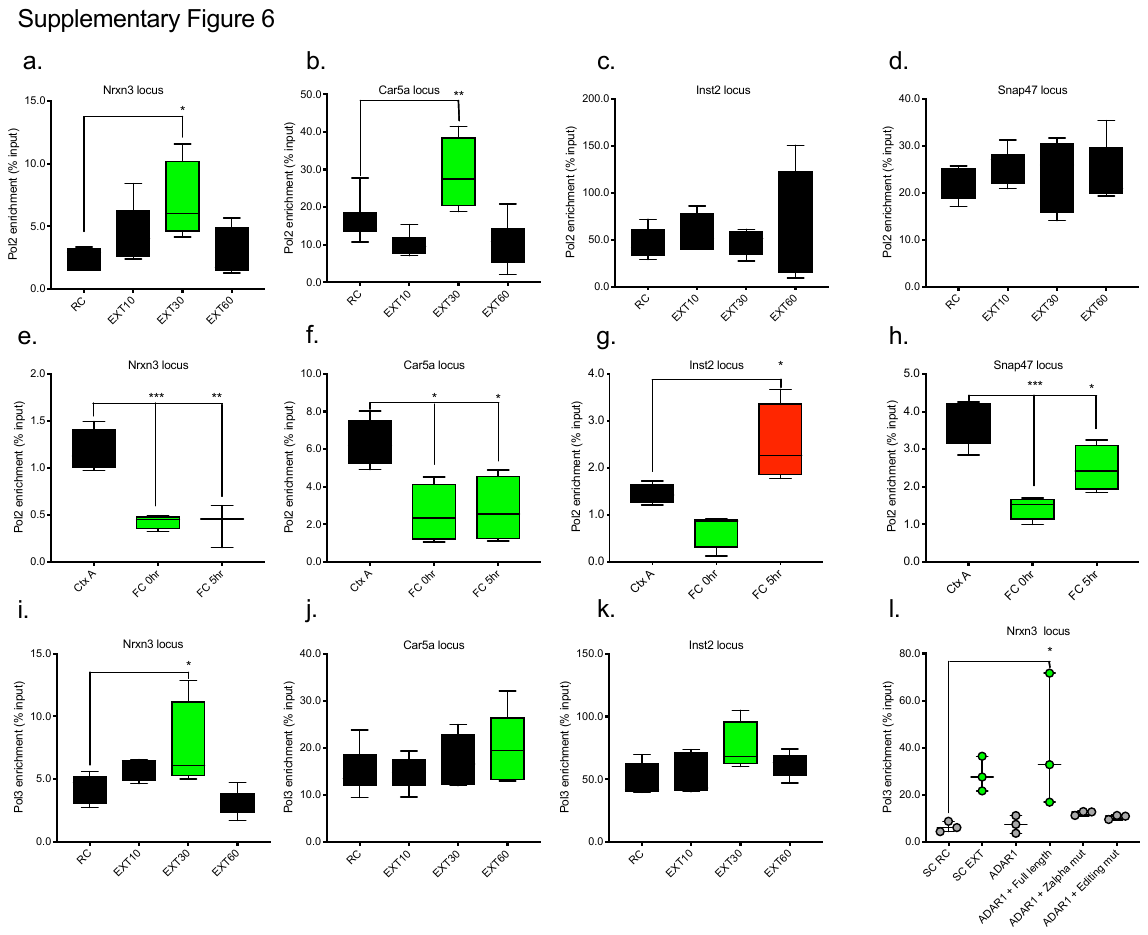

**Supplementary Figure 6. Polymerase II and III occupancy at ADAR1 hotspots**

Polymerase II (polII) enrichment significantly increased when comparing fear trained animals exposed to a novel context following fear conditioning (retention control: RC) to fear conditioned followed by 10CS (EXT 10), 30 CS (EXT 30), or 60CS extinction (EXT 60) at the **a,** Nrxn3 locus (ANOVA, F_3,20_ = 4.175, *p=0.0219, Dunnett’s posthoc tests: RC vs EXT 10 p = 0.2747 RC vs EXT 30 *p=0.0126, RC vs EXT 60 p=0.9852), and **b,** Car5a locus (ANOVA, F_3,20_ = 9.609 ***p=0.0004, Dunnett’s posthoc tests: RC vs EXT 10 p = 0.2106 RC vs EXT 30 **p=0.0088, RC vs EXT 60 p=0.2232). This change did not reach statistical significance at the **c**, Inst2 locus (ANOVA, F_3,16_ = 0.7212 p=0.5538) or (d) Snap47 locus (ANOVA, F_3,20_ = 0.2958 p=0.8279). For animals that were exposed to the fear conditioning context but not conditioned (Ctx A) compared to those fear conditioned and euthanized immediately after training (FC O hr), or 5 hr after (FC 5hr) there was a significant reduction in polII enrichement at the **e,** Nrxn3 locus (ANOVA, F_2,9_ = 21.85 ***p=0.0006, Dunnett’s posthoc tests: Ctx A vs FC 0hr ***p =0.0008, Ctx A vs FC 5hr **p = 0.0010), **f,** Car5a locus (ANOVA, F_2,9_ = 7.42 *p=0.0125, Dunnett’s posthoc tests: Ctx A vs FC 0hr *p =0.0136, *Ctx A vs FC 5hr p = 0.0187), **g,** Inst2 locus (ANOVA, F_2,9_ = 10.87 **p=0.0040, Dunnett’s posthoc tests: Ctx A vs FC 0hr p =0.1467, Ctx A vs FC 5hr *p = 0.0429), and the **h,** Snap47 locus (ANOVA, F_2,9_ = 18.52 ***p=0.0006, Dunnett’s posthoc tests: Ctx A vs FC 0hr ***p =0.0004, Ctx A vs FC 5hr p = 0.0160). Polymerase III (PolIII) showed a significant increase in binding at the **i,** Nrxn3 locus (ANOVA, F_3,16_ = 5.142, *p<0.05, Dunnett’s posthoc tests: RC vs EXT 10 p = 0.4125, RC vs EXT 30 *p=0.0457, RC vs EXT 60,) **j,** Car5a and **k,** Inst2 showed trends that did not reach significance. **l,** Additionally, Nrxn3 PolIII binding could be blocked by ADAR1 shRNA, and rescued by addition of full length ADAR1 (ANOVA, F_5,12_ = 3.922, *p<0.05 Fishers LSD: SC RC vs SC EXT 10 *p = 0.0440 SC RC vs ADAR1 EXT p = 0.9201, SC RC vs ADAR1 + Full length p = 0.0047, SC RC vs ADAR1 + Zalpha mut p = 0.5616, SC RC vs ADAR1 Editing mut).

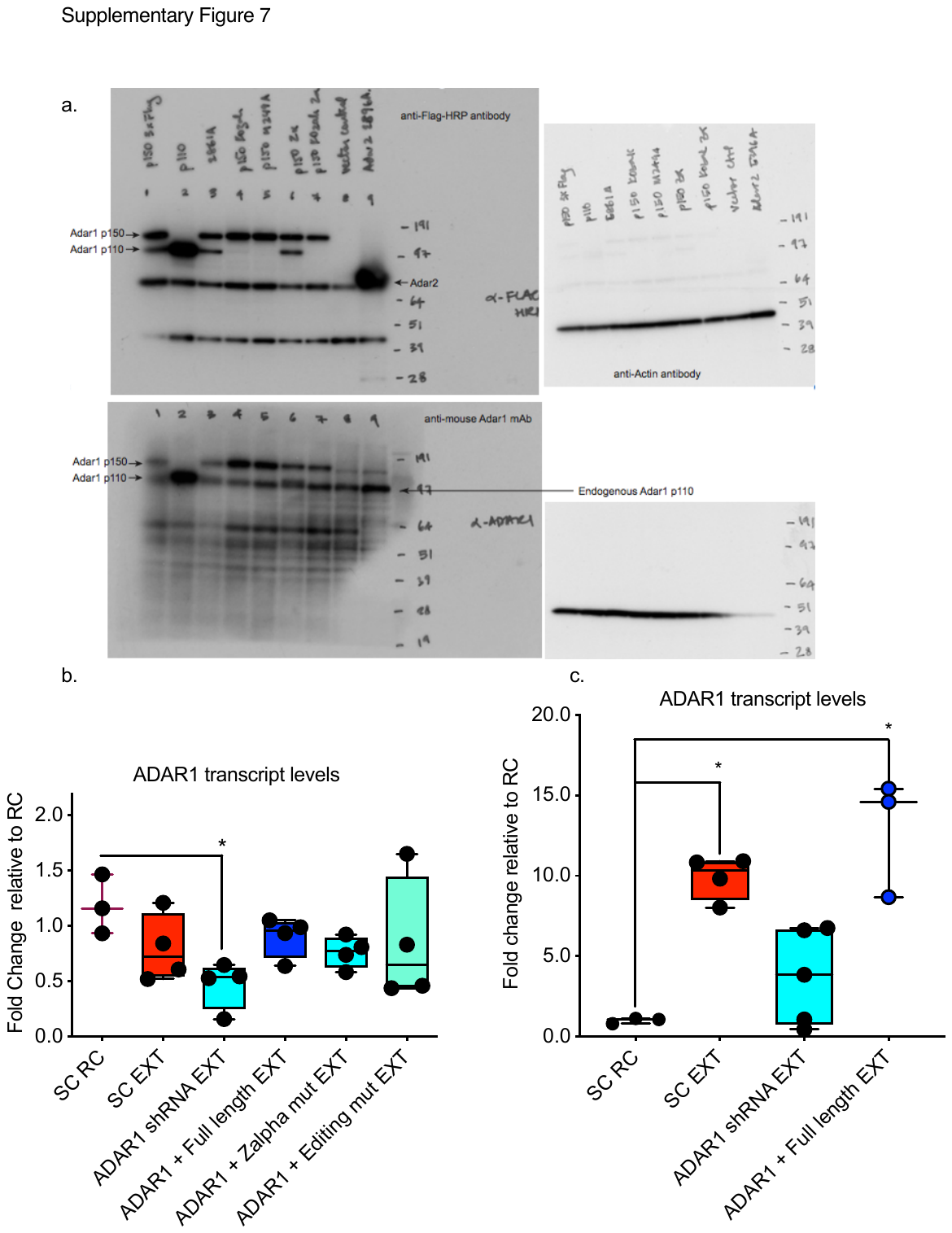

**Supplementary Figure 7. Validation of ADAR1 overexpression constructs**

**a,** Western blot representing constructs infected virus into murine Kusa4b10 stromal cell line:

1 = endogenous full length murine Adar1 construct - expressed both p110 and p150 (3xFlag with AgeI restriction site), 2 = endogenous p110 murine Adar1 construct - expressed only p110 (3xFlag with BclI restriction site), 3 = endogenous full length murine Adar1 E861A construct - expressed editing dead p110 and p150 (3xFlag with NsiI restriction site), 4 = endogenous murine Adar1 construct with optimized Kozak sequence - express p150 (3xFlag with NruI restriction site), 5 = endogenous murine Adar1 construct with optimized Kozak sequence and M249A (p110 initiation) mutation - express p150 (3xFlag with NruI restriction site), 6 = endogenous murine Adar1 construct with Za mutations N175A and Y179A - express p110 and Za mutant p150 (3xFlag with PmeI restriction site), 7 = endogenous murine Adar1 construct with Za mutations N175A and Y179A and optimised Kozak - express Za mutant p150 (3xFlag with PmeI restriction site), 8 = empty vector infected and puromycin selected cells, 9 = endogenous mAdar2 E396A editing deficient overexpression (3xFlag). **b,** Following the completion of training animals which had been exposed to fear conditioning and context B and treated with SC shRNA virus (SC RC) were compared to animals that underwent extinction training (EXT) and had either SC virus (SC EXT) ADAR1 shRNA (ADAR1 shRNA EXT), ADAR1 + full length ADAR1 (ADAR1 + FL EXT), ADAR1 virus + Zalpha mutant construct (ADAR1 + Zalpha EXT), or ADAR1 virus + editing mutant (ADAR1 + Edit mutant EXT). This demonstrated that ADAR1 was significantly depleted by ADAR1 shRNA (ANOVA F_5,17_ = 1.890, *p = 0.0414 SC RC vs. ADAR1 shRNA EXT, and that this could be restored by addition of any of the other domains **c,** Further activity dependent ADAR1 increase is fully restored by addition of full length domain, and blocked by ADAR1 shRNA (ANOVA, F_3,11_ = 1.507, ***p<0.001, Dunnet post hoc; **SC RC vs. SC EXT p = 0.0017, SC RC vs. ADAR1 shRNA EXT p = 0.3299, SC RC vs. ADAR1 + Full length EXT ***p = 0.0003).
